# Spatial Transcriptomics reveals a T cell-mediated microglial activation axis of neurodegeneration following immune checkpoint inhibition

**DOI:** 10.64898/2026.06.10.730982

**Authors:** Devyani Swami, Suhas Sureshchandra, Janaki Manoja Vinnakota, Robert Zeiser, Shivashankar Othy, Munjal M. Acharya

## Abstract

Immune checkpoint inhibitor (ICI) combinations that block cytotoxic T-lymphocyte–associated protein 4 (CTLA-4) and programmed cell death protein 1 (PD-1) signaling have revolutionized cancer care but also exert a range of immune-related adverse events (irAE) in various tissues, including the brain. Our understanding of the mechanisms of irAE in the brain is still evolving, and we recently demonstrated that ICI (blockade of CTLA-4 and PD-1) perturbs hippocampal-dependent memory function by derailing neuro-immune homeostasis and compromising synaptic integrity. However, the spatial patterns and the cell–type–specific molecular mechanisms underlying ICI-related brain dysfunction remain not well-defined. To address this gap, we performed spatial transcriptomic profiling of the hippocampal region using multiplexed error-robust fluorescence in situ hybridization (MERFISH) to map gene expression at single-cell resolution. By integrating spatial single-cell data with bulk RNA-seq, we define the distribution of microglia, astrocytes, synaptic, and neuroinflammatory markers, and determine how ICI reshapes hippocampal cellular composition in a syngeneic murine melanoma model. MERFISH revealed upregulation of microglial, astrocytic, oligodendrocytic, and T cell markers post-ICI treatment, revealing unique pathways driving neuroinflammation, synaptic function, and cellular signaling. Furthermore, immunofluorescence analysis of postmortem brains from patients treated with ICI corroborates our findings of ICI-related immune activation of microglia. Finally, using a conditional deletion model, we show that T cells are indispensable for ICI-driven microglial activation. Altogether, our study provides a high-resolution spatial framework for understanding irAEs in brain function and a T cell-microglia crosstalk axis as a driving mechanism of dysregulated neuro-immune homeostasis during ICI.

## INTRODUCTION

Immune checkpoint inhibitors (ICIs), targeting programmed cell death protein 1 (PD-1) and cytotoxic T-lymphocyte–associated protein 4 (CTLA-4), are transformative breakthroughs in modern oncology. ICIs have reshaped cancer treatment by restoring anti-tumor immunity, leading to major improvements in patient survival and overall quality of life (Mc Neil V et al., 2025). ICI therapies continue to expand rapidly and are translated across a wide range of advanced or metastatic cancers, including lung, renal, gastrointestinal, breast, and prostate cancers (Miller SR et al., 2024; Wei J et al.,2024). ICIs are administered either alone or in combination regimens to reinvigorate exhausted immune cells, particularly cytotoxic (CD8⁺) T cells, that play a central role in eliminating tumors (Buder-Bakhaya K et al., 2018; Hargadon KM et al., 2018). However, despite their therapeutic success, ICIs cause off-target effects and normal tissue toxicities (Darnell EP et al., 2020, Das Se et al., 2019). Cancer survivors, including individuals with melanoma and recipients of ICI, often report cognitive impairments (Goldberg SB et al., 2016; Rogiers A et al., 2020). Commonly reported neurologic events include encephalitis, headaches, seizures, fatigue, myasthenia gravis, Guillain-Barre syndrome, and peripheral neuropathy (Farina A et al., 2023; Haugh AM et al., 2020). For example, 41% of Ipilimumab (αCTLA-4)-treated survivors demonstrate objective neurocognitive dysfunction, even after 5-6 years post-treatment. In some cases, individuals are forced to discontinue employment (Rogiers A, et al., 2020a, (Rogiers A, et al., 2020b, Rogiers A, et al., 2023). Similarly, 32% of Pembrolizumab (αPD-1) survivors showed cognitive decline, accompanied by a reduced quality of life. The most concerning issue is that these deficits persist and worsen after the treatment completion, with increased emotional and cognitive impairment documented at 18 months post-cessation, independent of cancer recurrence (Rogiers A, et al., 2020a, (Rogiers A, et al., 2020b, (Rogiers A, et al., 2023). Up to 26% incidence of clinically significant cancer-related cognitive impairments (CRCI) in ICI-treated patients across lung cancer, GI cancers, and melanoma (Sayer M et al., 2025. The clinical burden extends to severe psychological sequelae, with cancer-related PTSD in 48% of survivors and 28% experiencing suicidal ideation (Masaki K et al., 2023), (Reynolds KL et al., 2019), (Ho MH et al., 2024), (Chan A et al., 2023), (Demos-Davies K et al., 2024). Moreover, ICI (αCTLA-4)-related neurotoxicity was associated with demyelination and CNS autoimmunity, and myelin-reactive CD4^+^ T cells were readily identifiable in ICI-treated patient brains((Cao Y et al., 2016). Despite increasing reports of concerning patterns of neurocognitive and psychosocial sequelae of ICI, the cellular and molecular mechanisms by which ICI orchestrates these neurodegenerative and neurocognitive complications remain understudied (Fleming B et al., 2023; Myers JS et al., 2023).

Using a murine model of ICI-responsive melanoma, we recently demonstrated that combinatorial ICI treatment (anti-PD-1 + anti-CTLA-4; Combi-ICI) effectively controlled tumor growth but led to a marked decline in learning, memory, and memory consolidation (Ifejeokwu OV et al., 2025). Combi-ICI induced glial dysfunction, characterized by microglial activation and astrogliosis, which amplified neuroinflammation, disrupted synaptic and neuronal integrity, and contributed to cognitive impairments. We also observed an increased pool of T cells and an upregulation of MHC-II⁺ myeloid cells in the brain and demonstrated that ICI exaggerates autoimmune encephalitis. Together, these neuroimmune perturbations significantly compromise hippocampal-dependent cognitive processing. However, the location and cell type-specific molecular mechanisms that drive ICI-mediated neuroinflammation and neuronal dysfunction remain poorly defined. In particular, how the spatial patterns of ICI-mediated changes in glial and immune cell states, and their interactions within the brain microenvironment. Therefore, there is a critical need for high-resolution, spatially resolved transcriptomic approaches to map ICI-induced molecular changes in the brain glial-immune network. Such methods would allow direct comparisons between melanoma-associated neuroinflammation and ICI-induced neuroimmune changes, thereby providing deeper insights into the transcriptomic and cellular pathways affected by ICI and, ultimately, offering potential mitigation strategies. Since these neuroimmune disturbances impair hippocampal-dependent learning, memory, and cognitive processing, vulnerability is likely shaped by the unique molecular signatures and spatial arrangement of its cell subtypes. Understanding this organizational landscape is essential for uncovering how ICI therapy disrupts hippocampal function and for guiding the development of targeted strategies that protect the most affected cell populations while preserving the anti-tumor effect of ICI.

This study establishes a spatial transcriptomic framework to dissect the cellular interactions and gene networks that mediate neurotoxic responses to ICI therapy. We mapped the spatial distribution and transcriptional states of microglia, astrocytes, and T cells, and the expression of key synaptic and neuroinflammatory signatures following Combi-ICI treatment. Our analyses revealed distinct changes in hippocampal cellular composition and gene expression in melanoma- and Combi-ICI–treated brains, highlighting pathological crosstalk among glial and inflammatory cell populations. To validate these findings in a translational context, we performed immunofluorescence staining using cell–type–specific markers on postmortem brains from ICI-treated patients with reported cognitive impairment. Although our transcriptional and cross-species translational analyses establish a mechanistic link between T-cell infiltration and microglial activation following ICI treatment, these results, more broadly, provide a high-resolution view of the molecular and cellular architecture underlying ICI-associated neurocognitive dysfunction.

## MATERIALS AND METHODS

### Animal model and injections

All animals used in this study were housed and cared for in accordance with National Institutes of Health (NIH) guidelines and protocols approved by the Institutional Animal Care and Use Committee (IACUC). Adult male wild-type (WT) mice aged 10–12 weeks were randomly assigned to one of the following experimental groups: (i) WT mice receiving isotype-matched control antibodies (intraperitoneal, i.p.); (ii) WT mice bearing D4M-3A.UV2 tumors and treated with isotype-matched control antibodies (i.p.); (iii) WT mice treated with immune checkpoint inhibitors (ICIs), including anti–CTLA-4 (1 mg per dose, three times weekly, i.p.) and anti-PD-1 (200 µg per dose, twice weekly, i.p.); and (iv) WT mice bearing D4M-3A.UV2 tumors treated with the same combination ICI regimen for three weeks.

For tumor induction, mice in the cancer groups received bilateral intradermal injections of 5 × 10 ^5^ syngeneic D4M-3A.UV2 melanoma cells. Tumors were allowed to engraft for one week prior to initiating ITC or ICI treatments. Following completion of the treatment regimen, mice were euthanized 72 hours after the final injection, and brains were harvested for downstream analyses.

### Tissue preparation, Transcriptomics, and Data processing

For Bulk RNA seq data, fresh frozen single hippocampi from N=5 brains /groups were excised from the mouse brain and were sent to Azenta Life science. Briefly, total RNA extraction, library preparation and sequencing using RNA samples were quantified using Qubit 2.0 Fluorometer (Life Technologies, Carlsbad, CA, USA) and RNA integrity was checked using Agilent TapeStation 4200 (Agilent Technologies, Palo Alto, CA, USA). The sequencing libraries were sequenced on the Illumina NovaSeq X Plus instrument The samples were sequenced using a 2×150bp Paired-End (PE) configuration, targeting approximately 30 M reads/sample. Further details are available in supplementary data. For differential expression gene analysis (DEGs) files, read counts were normalized to adjust for various factors, such as variations of sequencing yield between samples (software package DESeq2 v.1.16.1). For analysis, A Wald test analysis was used to test for changes in gene expression (software package DESeq2 (v.1.16.1)). Genes with adjusted p-values < 0.05 and an absolute log2-fold change > 1 were considered statistically significant differentially expressed genes. For subsequent analysis using DEG files, Benjamini Hochberg adjusted p values and statistical P values <0.001 were used for comparative analysis across the datasets.

For Spatial transcriptomics using MERFISH, whole brains were carefully harvested from mice following decapitation and immediately placed into tissue-embedding plastic molds. The prechilled OCT was poured onto tissue embedding plastic mold and later transferred into isopentane and liquid nitrogen bath until OCT completely solidifies and turns white. To ensure RNA integrity, at least the first 50 µm of tissue sections were discarded prior to collecting evaluable samples. Subsequently, 10 µm sections from the OCT-embedded tissue blocks were mounted onto slides and processed for probe hybridization and gel embedding according to the MERFISH protocol.

Prepared slides were assembled into the MERSCOPE Flow Chamber and loaded into the MERSCOPE instrument along with the appropriate imaging cartridge. Raw images were processed using the MERSCOPE software pipeline to generate spatial genomics output files ready for downstream analysis. These outputs included detected transcripts and their three-dimensional spatial coordinates, provided as cell metadata (CSV), experiment metadata (JSON), cell boundary files (HDF5), and MERSCOPE Visualizer files (VZG). Using the MERSCOPE Visualizer software, AnnData (HDF5) files were loaded and hippocampal regions of interest (ROIs) were selected. ROI-specific HDF5 files were subsequently exported and converted into RDS format in R for downstream differential gene expression (DEG) analysis.

### Analysis of MERFISH spatial transcripts data

We designed a MERFISH library of 300 genes relevant under three broad categories: (1) Excitatory markers; (2) Inhibitory markers (3) non- Neuronal markers. Cell type markers were based on the Gene panel-specific MERSCOPE Codebooks and were customized using MERSCOPE Gene Panel Design Software. Cell by gene matrix was imported from MERSCOPE visualizer software as HDF5 file and was loaded onto Seurat for downstream analysis.

Cells were first integrated across datasets using Seurat, followed by dimensionality reduction and graph-based clustering. Leiden clustering was performed on the integrated objects using 2,000 highly variable features at a resolution of 1.0, generating transcriptionally distinct clusters that were visualized using UMAPs. For each Leiden cluster, cluster-specific marker genes were identified (imported as a gene panel). Cell-type identities were assigned using a marker overlap–based scoring approach. Curated canonical marker gene sets were defined for major neural and immune cell populations, including astrocytes, microglia, oligodendrocytes, excitatory neurons, inhibitory neurons, T cells, B cells, and synaptic-function–enriched populations. Clusters meeting both criteria—canonical T-cell identity and complement gene enrichment—were labeled as T cells & Complement-enriched. Clusters expressing ≥2 T-cell markers were annotated as T cells. Clusters additionally expressing ≥2 complement pathway genes were labeled as Complement-enriched, clusters meeting both criteria were annotated as T cells (Complement-enriched). Leiden clusters within each cell type were relabeled with simplified identifiers (e.g., A1–An for astrocytes, O1–On for oligodendrocytes, T1–Tn for T cells, and M1–Mn for microglia), without altering the underlying clustering structure. These relabeled cluster identities were stored as metadata and used exclusively for visualization. To assess condition-specific distributions of cell type–specific clusters, UMAP embeddings were visualized using DimPlot and split by experimental condition. Importantly, the same UMAP coordinate system was used across all split panels, enabling direct comparison of cluster abundance and spatial organization across conditions. Volcano plot across the cell types and across the dataset was generated using EnhancedVolcano package, with log₂ fold change (avg_log2FC) plotted against adjusted p-value (–log₁₀ scale). Upregulated and downregulated genes were visualized simultaneously, enabling direct comparison of transcriptional changes between conditions. Overlapping gene sets exhibiting opposite regulation between groups were identified using set intersection analysis.

Gene ontology

Functional enrichment analysis of differentially expressed genes (DEGs) for selected cell types was performed using Metascape. Enriched Gene Ontology (GO) biological processes and pathway terms were identified using default Metascape parameters. Bubble plots were generated in R using ggplot2 to visualize selected enriched GO terms across experimental conditions. Enrichment patterns were compared across experimental groups (Control, Control + ICI, Melanoma, and Melanoma + ICI) and microglial subclusters. Bubble color represents the scaled average expression of genes associated with each term, while bubble size indicates the percentage of cells expressing the corresponding genes within each cluster.

### Dual-immunofluorescence staining, confocal microscopy, and 3D algorithm-based volumetric quantification

*Mouse brains.* Free floating mouse brain sections (N=4-8 brains/group, 2-3 sections/ brain) with visible hippocampi and tumors (N=3 tumors, 3 sections/tumor) were chosen for immunohistochemistry (IHC). Primary antibodies include Rat anti Mouse CD68 (1:500, Bio-Rad, Cat# MCA1957; RRID: AB_322219), Rabbit anti IBA1 (1:500, Wako, Cat# 019-19741; RRID: AB_839504)

For staining, tissue sections were first washed 3 times in phosphate-buffered saline (1X PBS, 100mM, pH 7.4, FisherSci) and, when necessary, antigen retrieval was performed using citrate buffer incubation (10mM, pH 6.0 with 0.05% Tween-20, Sigma) 70°C for 45 minutes. Tissues were then incubated in borate buffer (pH 8.5, 100mM, Sigma, Cat. B0394) for 10 minutes, followed by three 5-minute washes in 1X PBS. Tissues were then blocked in serum (10% normal goat or donkey, NGS or NDS) in PBS with 0.01% Triton X-100 (NDS, Jackson ImmunoResearch Labs Cat# 017-000-121, RRID: AB_2337258; NGS, Jackson ImmunoResearch Labs Cat# 005-000-121, RRID:AB_2336990). Following an hour of incubation in the blocking solution, the tissues were incubated in primary antibodies. Primary antibodies were prepared in 0.01% Triton X-100 (Sigma) 3% NGS or NDS in PBS and incubated for at least 12 hours at 4°C in a shaker incubator (Enviro-Genie Scientific). The tissues were washed in 1X PBS (3 times) and then incubated in the secondary antibodies for one hour. Finally, sections were counterstained with DAPI nuclear dye in PBS for 15 min (1 µmol/L, FisherSci, Cat# D1306, RRID:AB_2629482). Finally, sections were washed in PBS and then mounted on superfrost slides (FisherSci) using Antifade mounting medium (VectaShield Cat# H-1000-10).

*Human Brains.* FFPE sections from human cortical tissues (frontal cortex) isolated from immune checkpoint inhibitor (ICI)-treated or non-tumor control brains post-mortem were gifts from Department of Nuclear Medicine, LMU Munich, Medical Center-University of Frieburg. The use of the human brain samples for this study was approved by the Institutional Biosafety committee of University of California, Irvine. Details about the patient treatment are available in the **Suppl. Table 6 “C:\Users\swami\Downloads\Supplemental Table Figures.docx“**. Primary antibodies include Rabbit anti-CD16 /Fcgr3 (1: 500, Fortis Life Science, Cat# A700-163-T, BLR163J), Mouse mAb Trem2 (1:500, Cell signaling Technology, Cat# 29715T, E4F5G) GFAP, Mouse Alpha-1-Antichymotrypsin / anti-SERPINA3 (1:500, Mybiosource.com, Cat# MBS439876, SERPINA3), Mouse anti-Human SOX10 (1:500, Mybiosource.com, Cat# MBS439095, So× 10), Rabbit Anti-GFAP (1:400, Biolegend, Cat# 840001, PRB-571C), Mouse Myelin Basic protein (MBP) mAb (1:500, Origene, Cat# SM1453S, Clone ID: 22)

For staining, formalin-fixed, paraffin-embedded (FFPE) human brain tissue sections were processed using a standard immunofluorescence protocol. Frontal cortex tissue from three patients with different malignancies who had received ICI treatment (anti-PD-1, or Combi-ICI), along with three non-tumor, untreated control patients, was used. Four tissue sections from each patient brain sample were stained and quantified. Briefly, Slides were first deparaffinized by incubation in Histo-Clear for 20 min, followed by sequential rehydration in 100%, 90%, and 70% ethanol for 3 min each, and a final rinse in distilled water for 1 min. Antigen retrieval was performed by immersing slides in antigen retrieval solution (Citrate buffer solution) within a glass chamber and heating in a microwave for 4 min (optimized), ensuring boiling for at least 3 min, this heating–cooling cycle was repeated once, after which slides were allowed to cool completely. Slides were then gently wiped to remove excess retrieval solution, washed under running distilled water for 10 min, and equilibrated in 1× TBST for two washes of 5 min each. For blocking, sections were incubated in a humidified chamber with blocking buffer containing 10% goat serum and 5% BSA prepared in 1× TBST for 1 h 30 min at room temperature. Primary antibodies were diluted 1:400 in incubation buffer (1% BSA, 1% goat serum in 1× TBST) and applied overnight at 4 °C. Slides were subsequently washed three times in 1× TBST for 5 min each and incubated with the appropriate secondary antibodies diluted 1:500 in the same incubation buffer for 60 min at room temperature. Following secondary incubation, slides were rinsed in distilled water for 1 min and washed three times in 1× TBST for 10 min each. Nuclear staining was performed using in-house 10× DAPI solution for 10 min (1 µmol/L, FisherSci, Cat# D1306, RRID:AB_2629482), followed by a final wash in 1× TBST for 10 min. Sections were mounted using VECTASHIELD mounting medium (VectaShield Cat# H-1000-10) and stored protected from light until imaging. We randomly selected 3 to 4 regions of interest (ROIs) from each sections from the non-cancer control and ICI-treated patient’s brains for the volumetric quantification of immunoreactivity of each glial marker.

### Conditional T cell depletion and Combi-ICI treatment

T-DTR mice (Cd4-Cre^+/-^ DTR^+/-^) were generated by crossing of CD4-Cre (JAX# 017336: Tg(Cd4-cre)1Cwi/Bflu) mice with B6-iDTR (JAX#007900: C57BL/6-Gt(ROSA)26Sortm1(HBEGF)Awai/J). Because of expression of Cd4 during the double positive (CD4+ CD8+) stage of thymic selection, T-DTR mice express diphtheria toxin receptor (DTR) selectively in both CD4⁺ and CD8⁺ T lymphocytes. To achieve robust T cell depletion, T-DTR mice were injected with diphtheria toxin DT (100 ng, once daily for 2 days) at 2 days before or 7 days after ICI treatment to determine the impact of T cell ablation on microglial activation state. Dual immunofluorescence staining for microglial activation (CD68-IBA1, **Fig. 8**) was conducted in brains collected at 72 hours after either ICI or DT treatment

### Microscopy and 3D Algorithm-based Volumetric Quantification

Laser-scanning confocal microscope (Nikon Eclipse Ti2 AX) was used to image the immunostained brain sections at high resolution (2048 xy pixels, 24 µm thick z-stacks with 0.5 µm per z-stack). Acquisition of confocal z stacks, deconvolution, and 3D algorithm-based volumetric quantification of immunofluorescent punta was carried out as described in detail previously (10). 3D volume surfaces were created and quantified for each antigen of interest. Dual IHC stains such as Fcgr3/Trem2, Sox10/MBP, GFAP/Serpin3n, and co-localization volumetric between the 3D surfaces of the two markers were determined and automatedly quantified by Imaris (Oxford instruments). All images for each IHC stain were uniformly applied with the same parameters to obtain unbiased analyses

### Statistical analysis

All data are expressed as the mean ± SEM. Statistical analyses of immunohistochemical data were conducted using two-way ANOVA and t-test (non-parametric) (GraphPad Prism, v8.0). For gene expression analysis, the log2 fold change (magnitude of expression difference between conditions), and the –log10 of the adjusted P-value (statistical significance after multiple testing correction, Benjamini-Hochberg FDR) were used.

## RESULTS

### Melanoma-modulated neuroinflammatory signaling persists after immune checkpoint inhibition

To investigate global transcriptional alterations underlying neuroimmune dysregulation following ICI therapy in melanoma-bearing mice (Ifejeokwu et al., 2025), we performed bulk RNA sequencing on hippocampal tissue collected 3 days after completion of ITC or Combi-ICI (αCTLA-4 + α PD1) treatments (**Fig. 1A**). Differential expression analysis was conducted across three pairwise contrasts: Control vs. Melanoma, Melanoma vs. Melanoma + ICI, and Control + ICI vs. Melanoma + ICI (**Fig. 1B**). We observed that melanoma-bearing mice exhibited a pronounced transcriptional activation signature relative to controls, consistent with tumor-driven neuroinflammatory responses. In contrast, ICI-treated tumor-bearing mice showed widespread downregulation of genes compared with the Melanoma group, suggesting that ICI may partially attenuate tumor-induced immune activation. Notably, the Melanoma + ICI group still displayed a substantial number of upregulated genes compared with the Control + ICI group, indicating that tumor-associated transcriptional activation persists in the hippocampus despite ICI therapy. A complementary three-way Venn diagram further revealed unique patterns (**Fig. 1C)**. The Control vs. Melanoma comparison contained seven uniquely upregulated genes *Adora2a, Fosb, Htr1b, Lcn2, Sult1a1, Fmo2, Acer2* with an additional three genes shared with the Control + ICI vs. Melanoma + ICI group *Adora2a, Cd4, Lcn2*. Conversely, the Melanoma + ICI comparison contained four uniquely upregulated genes *Htr7, Nog, Glra3* highlighting tumor-induced transcriptional changes that remain present post-ICI treatment.

**Figure 1.**
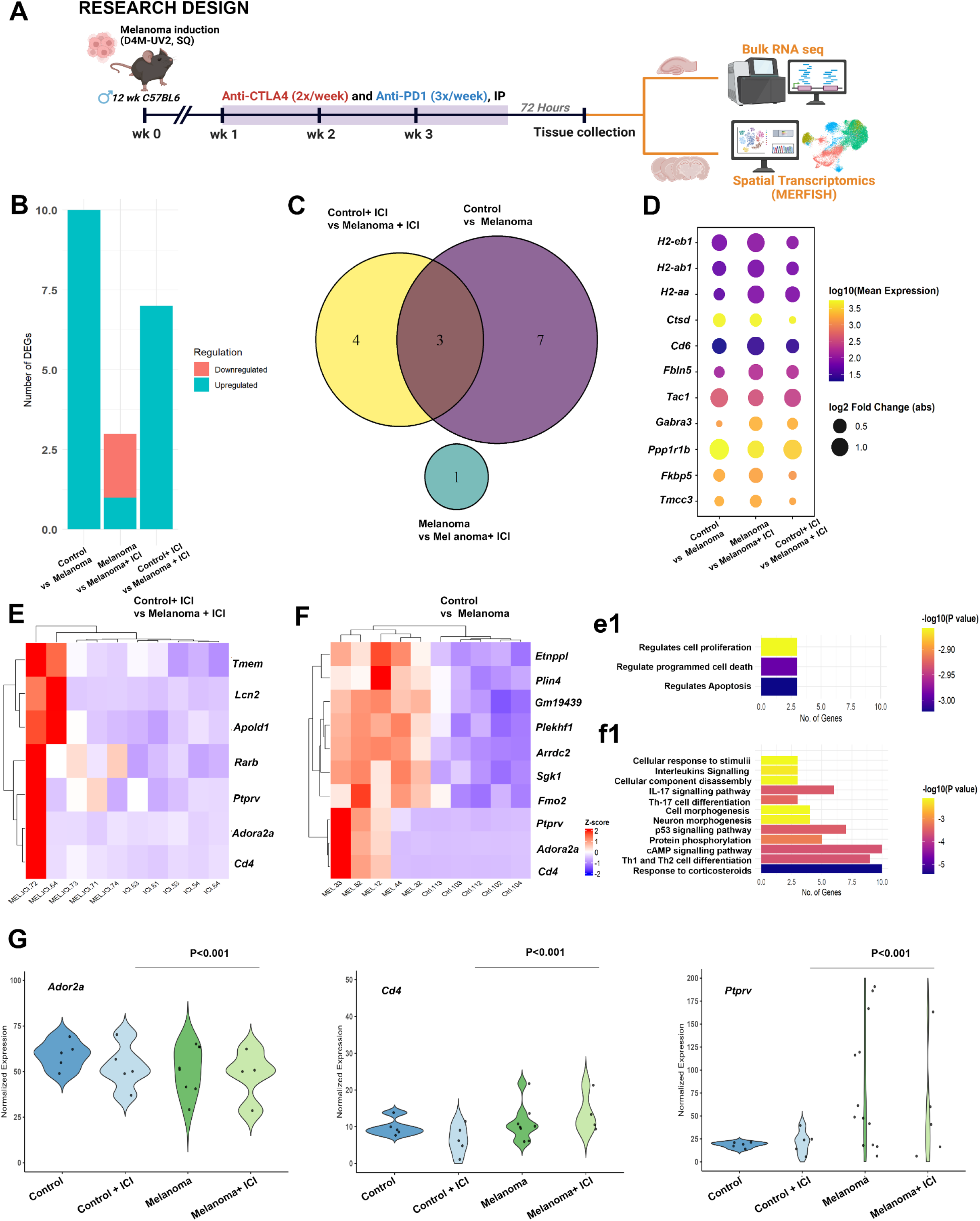
Bulk RNA sequencing of hippocampus post-ICI treatment. (**A**) Experimental design: Two-month-old C57Bl/6 WT male mice received either sham treatment or melanoma induction (D4M-UV2 cells; 1 × 10⁶ cells/animal, subcutaneous bilateral injections). Seven days later, mice were treated with either combination immune checkpoint inhibitors (ICI; anti-CTLA4: 1000 μg, twice weekly; anti-PD-1: 200 μg, thrice weekly; intraperitoneally) or vehicle for three weeks. On day 28, hippocampal tissue / whole tissue was dissected for the Bulk RNA seq and MERFISH, respectively.. (B) Stacked bar plot showing the number of differentially expressed genes (DEGs) in each group. (**C**) Venn diagram with corresponding functional enrichment analysis of genes upregulated across the groups. Statistically significant analytes among Control, Melanoma, Control ICI, and Melanoma ICI groups are indicated (#P<0.1). (**D**) Bubble plot depicting dysregulated cell-type markers in the melanoma ICI group. Bubble size reflects fold change, while color indicates gene expression levels. **(E-F)** Heatmaps and corresponding enriched pathways showing fold changes and expression patterns of upregulated genes across Control, Melanoma, and Melanoma ICI groups. (e1-f1) Functional enrichment analysis of upregulated genes from Control vs Melanoma and Control ICI vs Melanoma ICI was performed using Metascape based on their predicted target genes. Here, the x-axis represents the number of genes from the dataset associated with each enriched pathway, the y-axis lists the corresponding biological processes or pathways identified through enrichment analysis. The color scale indicates the statistical significance of enrichment expressed as –log₁₀(P value). **(G)** Violin plot showing commonly upregulated genes in both the Control + ICI vs. Melanoma + ICI and Control vs. Melanoma comparisons. These genes showed strong upregulation and high expression levels across both conditions. Data are derived from N = 4 mice per group.

Furthermore, we compared the selected gene transcripts across all experimental groups. Notably, *Ctsd* (a microglial lysosomal enzyme involved in antigen-processing pathways) was highly expressed in both the Melanoma and Melanoma + ICI groups **(Fig. 1D)**. In addition, the MHC-II–associated microglial genes *H2-eb1, H2-ab1,* and *H2-aa* were moderately expressed but showed pronounced upregulation in the Melanoma + ICI group, suggesting a shift of microglia toward an immune-activated state. Similarly, *Darpp-32* (a neural signaling and plasticity-related gene) was strongly upregulated in the Melanoma + ICI group, indicating that ICI therapy may modulate neural plasticity.

Next, we examined transcriptional changes associated with melanoma and ICI treatment. In the Melanoma + ICI group, the heatmap revealed strong upregulation of several genes (**Fig. 1E)**, including *Tmem252, Lcn2, Apold1, Rarb,* Ptprv*, Adora2a, and Cd4*, compared to the Control + ICI group. Pathway enrichment analysis indicated activation of processes such as negative regulation of cell population proliferation and programmed cell death (apoptosis), suggesting that ICI treatment further enhances neuroimmune activation in the presence of melanoma (**Fig. 1e1**). Similarly, in the Melanoma group, a high fold-change and statistically significant upregulation was observed in genes including Ptprv*, Adora2a, Cd4, Etnppl, Plin4, Fmo2, Sgk1, Plekhf1, and Arrdc2* compared to the Control group (**Fig. 1F)**. DEG-associated pathways in this comparison included T helper cell activation, IL-17 signaling, and broader interleukin-mediated immune pathways, reflecting strong immune activation and T cell-associated transcriptional responses in melanoma-bearing mice (**Fig. 1f1**).

As shown in the violin plot (**Fig. 1G)**, three genes (*Adora2a, Cd4, and Ptprv*) were significantly upregulated (P < 0.001) in both the Control + ICI vs. Melanoma + ICI and Control vs. Melanoma comparisons (Fig. 1F). These genes exhibited strong upregulation and high expression levels across both conditions. *Adora2a* (adenosine A2A receptor) is known to mediate microglial activation (Biswas K., 2023), while *CD4* is a canonical marker of T cell activation. PTPRV, a tyrosine phosphatase, and act as a potent inhibitor of cell proliferation when highly expressed (Doumont G. et al., 2005). Together, these findings indicate that the commonly upregulated genes form a core melanoma-driven transcriptional program that persists post-ICI treatment. Importantly, neuroinflammation triggered by melanoma burden was not reversed by ICI therapy and sustained in the hippocampal microenvironment post-ICI treatment.

### Spatial transcriptomic profiling reveals region-specific cellular organization and transcriptional diversity across hippocampal subfields

To investigate the cell-type-specific and spatial features of these alterations across the dataset, spatial transcriptomic profiling was performed on hippocampal sections from four experimental groups of adult male wild-type (WT) mice: Control, Control + ICI, Melanoma, and Melanoma + ICI **(Fig. 1A)**. MERFISH-based spatial transcriptomic analysis of the hippocampus resolved distinct neuronal and non-neuronal cell populations across major subregions, including CA1, CA3, dentate gyrus, and subiculum (Fig. 2A). Brains from three mice per group were collected, yielding ∼3,000–6,000 cells per sample. High-quality data from hippocampal cells was obtained across all experimental groups ;10,500 from the Control, 14,100 from the Control + ICI, 10,000 from the Melanoma, and 12,500 from the Melanoma + ICI groups, across three independent biological replicates per group. Unsupervised clustering revealed the major neuronal and glial cell populations, including excitatory neurons, inhibitory neurons, and non-neuronal cells, establishing a spatially resolved atlas for evaluating treatment-associated changes. To assign biological identities to the transcriptionally defined clusters, we combined cluster-specific marker genes with a curated gene-class annotation table **(Suppl. Table S1) “C:\Users\swami\Downloads\Supplemental Table Figures.docx”.** To visualize the distribution of excitatory, inhibitory, and non-neuronal cell populations across hippocampal subregions, we generated Uniform Manifold Approximation and Projection (UMAP) embeddings for each region (CA1, CA3, dentate gyrus, and subiculum, **Fig. 2A)**. The regional IDs corresponding to these subfields were obtained from the Allen Brain Atlas (e.g., CA1 = 382, CA3 = 400, DG = 402, Subiculum = 418). Each point in the UMAP represents a single cell (or voxel-averaged expression profile), colored by its annotated cell type, including excitatory, inhibitory, or non-neuronal. The resulting plots highlight region-specific segregation and transcriptional diversity within the hippocampus. The resulting visualizations highlight robust region-specific segregation and transcriptional diversity across hippocampal subfields, reflecting both anatomical and molecular specialization within the hippocampus. To visualize the proportional distribution of excitatory, inhibitory, and non-neuronal cell populations across hippocampal subregions (CA1, CA3, dentate gyrus, and subiculum), we created a donut plots **(Fig. 2a1).** Together, non-neuronal cells comprise the largest population in the CA1 region of the hippocampus, followed by all excitatory neurons and then the inhibitory neurons. Furthermore, using ROI-based annotation of hippocampus sections, we assigned MERFISH-profiled cells to CA1, CA3, DG, and Subiculum and quantified the distribution of non-neuronal cell classes across these hippocampal subregions. Cell identities were derived from cluster-level marker matching, after which regional cell counts were visualized using stacked horizontal bar plots. Among the non-neuronal populations captured by ROI-based MERFISH annotation, CA1 contained the highest overall number of cells, narrowly followed by DG, with lower counts in Subiculum and CA3 (**Suppl. Fig. S1**) “C:\Users\swami\Downloads\Supplemental Table Figures.docx”. Across all four hippocampal subregions, oligodendrocytes and microglia were consistently detected at appreciable levels, with astrocytes showing relative enrichment in DG. In contrast, T-cell and T-cell/Complement-enriched populations were present at lower frequencies across regions. Overall, these data support region-specific variation in non-neuronal cellular composition across the ROI-defined hippocampal subregions

**Figure 2.**
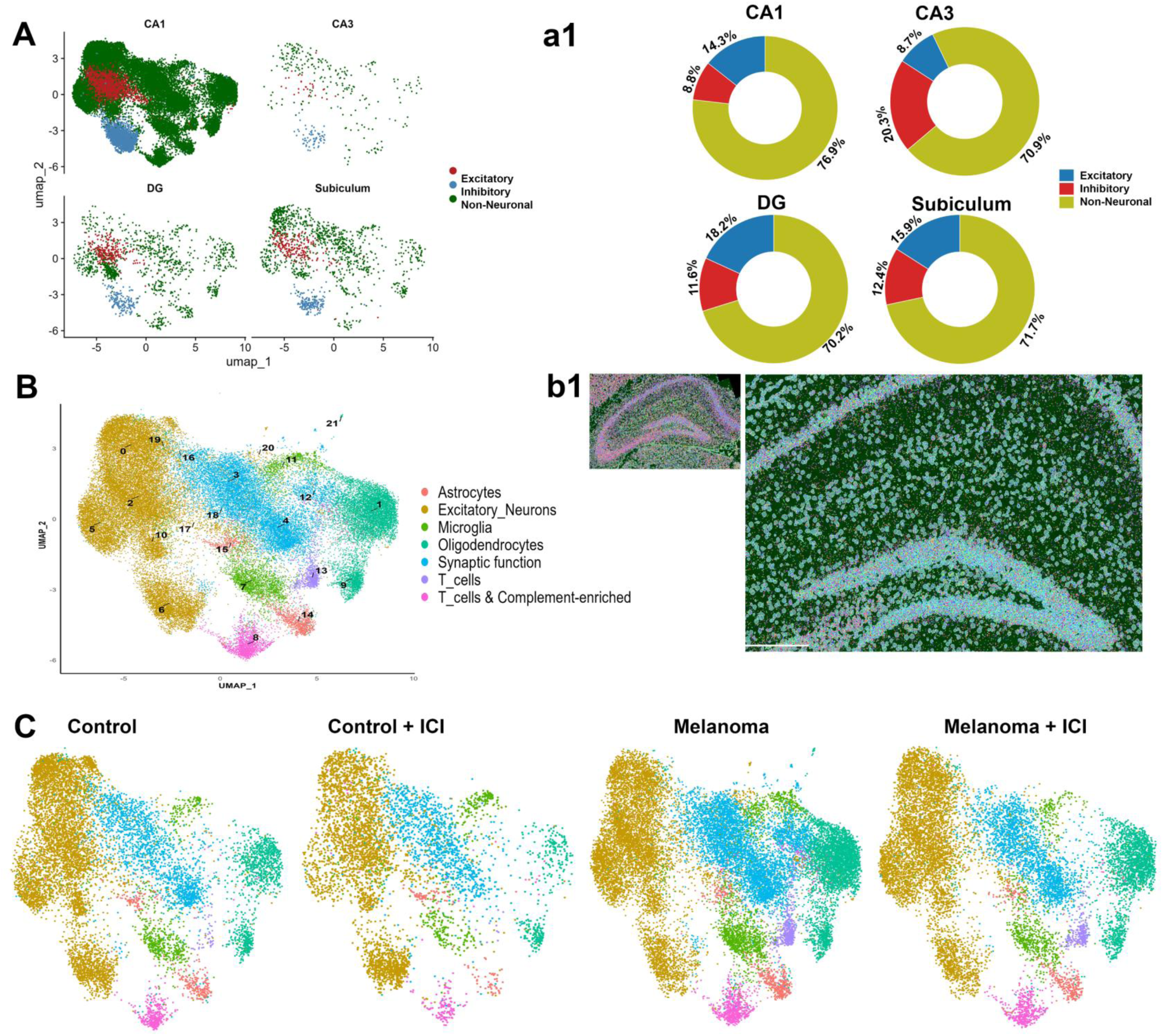
Spatial transcriptomic analysis of hippocampus post-ICI treatment. **(A)** Uniform Manifold Approximation and Projection (UMAP) embeddings for each region (CA1, CA3, dentate gyrus, and subiculum). (a1) Donut plots showing the proportional distribution of excitatory, inhibitory, and non-neuronal cell populations across hippocampal subregions (CA1, CA3, dentate gyrus, and subiculum) **(B)** UMAP visualization shows 21 clusters representing over 7 annotated cell types, following integration across all the experimental groups. (b1) Visualization of nuclei, segmented cell boundaries, and spatially mapped transcripts via MERSCOPE software **(C)** UMAP projections stratified by the experimental group dataset revealed clear differences among Control, Control + ICI, Melanoma, and Melanoma+ ICI groups Data are derived from N = 3 mice per group.

To further evaluate our detection accuracy, we first performed an integrated analysis of the MERFISH data. The UMAP visualization shows 21 clusters representing over 7 annotated cell types following integration across all experimental groups (Fig. **2C, c1**). The custom-made MERFISH gene library comprised 300 genes, including cell-type and functional markers for microglia, astrocytes, oligodendrocytes, immune cells, neurons, synaptic function, and CNS complement cascade-enriched markers. UMAP projections stratified by the experimental group dataset revealed clear differences among the Control, Control + ICI, Melanoma, and Melanoma + ICI groups (**Fig. 2C**). In particular, oligodendrocytes, astrocytes, T cells, microglia, and synaptic function-associated gene sets showed marked variation across conditions. For example, oligodendrocyte proportions were dramatically reduced in the Control + ICI group (6.38%) compared to the Melanoma and Melanoma + ICI groups (19.2% and 17.4%, respectively). Conversely, synaptic function-associated gene markers were substantially enriched in the Melanoma group (26.18%) relative to the Control and ICI-treated groups (Con + ICI and Melanoma + ICI). Similarly, T cells were more abundant in Melanoma (3.14%) and Melanoma + ICI (2.46%) than in animals without cancer (**Suppl. Table T1) “C:\Users\swami\Downloads\Supplemental Table Figures.docx”.**

### Microglial activation, a key driver of pro-inflammatory dysregulation, following ICI therapy

To investigate microglial dysregulation that may contribute to combined ICI–mediated synaptic loss, as previously reported (Ifejeokwu et al., 2025), (Fig. 3A). Microglia were identified using canonical gene markers and re-clustered with Seurat’s graph-based clustering, yielding multiple transcriptionally distinct were subsequently relabeled as M1–M16 for clarity **(Suppl. Fig. S2a**) “C:\Users\swami\Downloads\Supplemental Table Figures.docx”. To assess relative distributions of these subpopulations under different experimental conditions, UMAP plots were split by the experimental groups (Control, Control + ICI, Melanoma, and Melanoma + ICI, **Fig. 3A)**. Each dot represents a single microglial cell, and cells are colored according to their assigned cluster identity (e.g., M1–M16). This coloring highlights transcriptionally distinct microglial subpopulations and enables direct visual comparison of how the relative abundance and distribution of these clusters vary across experimental conditions Clustering of the microglial subset using the integrated microglial Seurat object at resolution 1.0 generated 16 microglial clusters, reflecting substantial transcriptional diversity. This resolution was selected because it captured meaningful microglial subpopulations while avoiding over fragmentation and maintaining cluster stability. The biological relevance of these clusters was further supported by heatmap analysis of microglial marker genes, which revealed cluster-specific gene expression profile across homeostatic and activation-associated microglial transcripts (**Suppl. Fig. S2b**) “C:\Users\swami\Downloads\Supplemental Table Figures.docx”.

**Figure 3.**
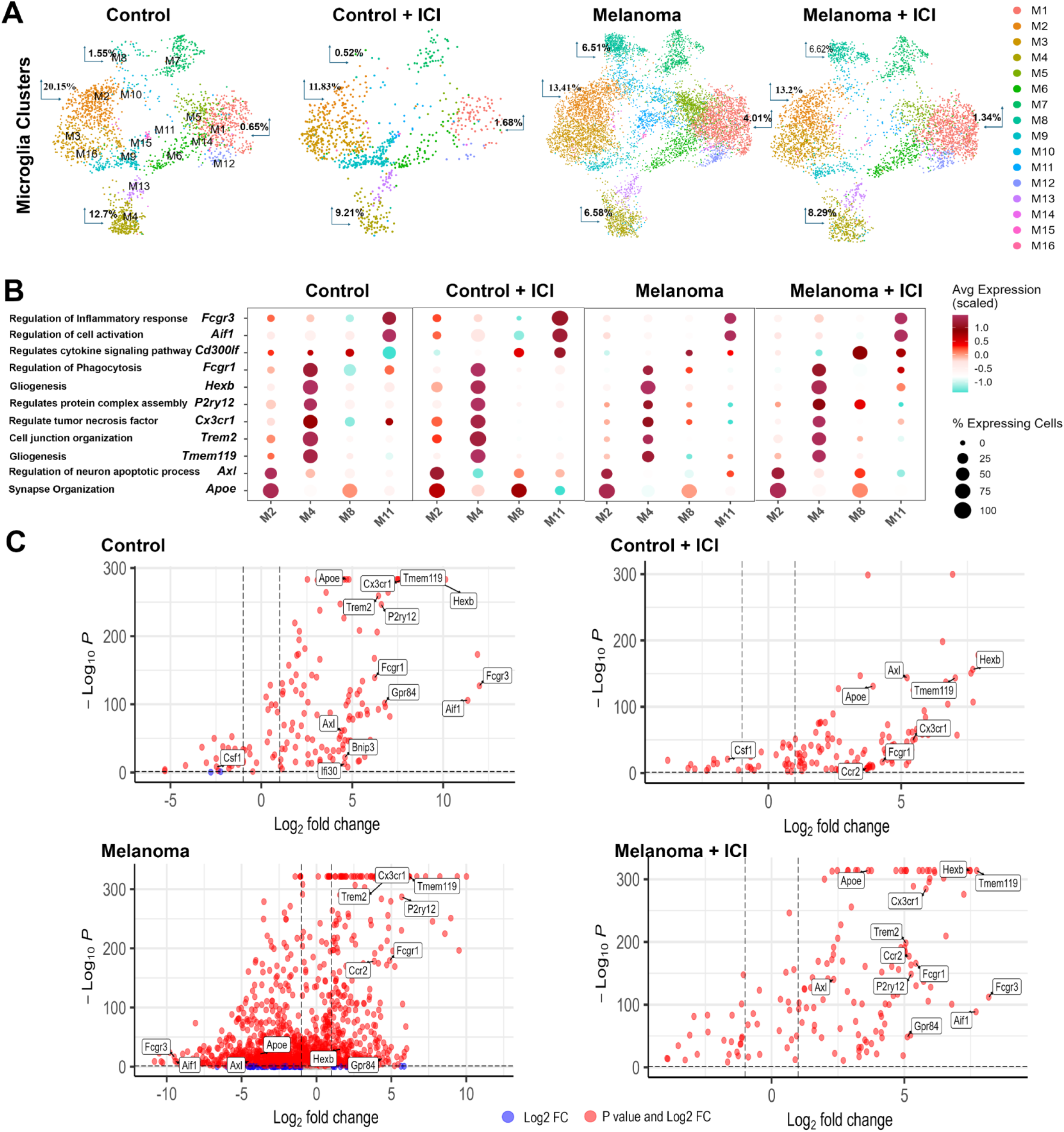
Gene expression analysis of the microglial cluster. **(A)** Split UMAPs display gene clusters across Control, Control +ICI, Melanoma, and Melanoma +ICI group datasets. **(B)** Bubble plots depicting the distribution and expression of genes within clusters. The size of the bubble represents the percentage of cells expressing the specific gene. The bubble color indicates the average expression level of that gene within the corresponding cluster. **(C)** Volcano plots depicting differential gene expression across the experimental group, with each point representing a gene. The X-axis corresponds to the log2 fold change (magnitude of expression difference between conditions), while the y-axis corresponds to the –log10 of the adjusted P-value (statistical significance after multiple testing correction, Benjamini-Hochberg FDR). Downregulated genes are further to the left, while upregulated genes are further to the right. Higher –log_10_*P* values indicated higher statistically significant gene expression. Significance thresholds used were: adjusted P < 0.05 and log2 fold change > 1. Red: Strongly upregulated, high significance. Blue: Strongly downregulated, high significance Data are derived from N = 3 mice per group.

Pairwise Fisher’s exact tests demonstrated that microglial clusters M2, M4, M8, and M11 exhibited significant alterations across experimental conditions **(Suppl. Table S2) “C:\Users\swami\Downloads\Supplemental Table Figures.docx“**. Clusters M2, M4, M8, and M11 were markedly enriched in Melanoma and Melanoma + ICI compared with Control, with the strongest effects observed for clusters M4, M8, and M11. These clusters showed highly significant differences across nearly all melanoma-related comparisons. In contrast, Control vs. Control + ICI comparisons revealed more modest but still significant shifts for clusters M2, M4, M8, and M11 (with a slight change in M7), indicating that ICI alone alters microglial composition independent of tumor.

To quantify microglial cluster composition (M1–M16) across groups, we calculated the per-sample proportion of cells assigned to each cluster and summarized these distributions using 100% stacked bar plots **(Suppl. Table S2) “C:\Users\swami\Downloads\Supplemental Table Figures.docx“**. Further characterization of the functional identities of each cluster highlighted their defining marker genes and expression patterns across datasets **(Fig. 3B)**. Briefly, in the Control + ICI group, clusters M4 and M11 contained a higher proportion of cells, with ∼100% of cells expressing *Aif1* (a microglial activation marker; De Leon-Oliv et al., 2023) and *Fcgr3* (a regulator of microglial polarization; Chauhan et al., 2017). The concurrent upregulation of *Aif1* and *Fcgr3* suggests that these clusters shift toward an activated, pro-inflammatory state, consistent with Fcgr3’s known role in modulating inflammatory responses under ICI treatment. Approximately 75% of genes expressed in cluster M4 across the Control, Control + ICI, and Melanoma + ICI groups corresponded to canonical homeostatic microglial markers, including *Trem2, Tmem119, P2ry12, Cx3cr1, and Hexb*. The relative loss of this homeostatic signature in the Melanoma group indicates a disruption of baseline microglial maintenance functions. Similarly, cluster M8, which expanded in both vehicle- and ICI-treated melanoma groups compared with Control, contained a high proportion (∼75%) of cells expressing *Apoe*. In contrast, the larger M8 cluster in the Melanoma + ICI group was enriched for *Cd300lf,* suggesting lower neuroinflammatory activity (Lu Z, et al., 2024) and a possible role for microglia in promoting neurological recovery following ICI therapy.

To investigate gene-level insights complementary to the cluster-based observations, we next sought to identify individual genes that were significantly altered across datasets. We performed differential expression analysis of microglial populations and visualized the results using volcano plots. As shown in the figure **(Fig. 3C)**, the key homeostatic microglia marker Hexb was strongly upregulated in the Control, Control + ICI, and Melanoma + ICI groups, but weakly upregulated in the Melanoma group. This pattern suggests that in the Melanoma group, microglial homeostatic programs are disrupted, whereas ICI partially restores or preserves these features. Similarly, *Gpr84*, expressed by activated microglia, was upregulated under pathological conditions, consistent with previous reports (Audoy-Rémus et al., 2015). In line with these findings, our results demonstrate that *Gpr84* was significantly upregulated in both the Melanoma and Melanoma + ICI groups, supporting its role as an inflammatory amplifier of microglial pathophysiology. Overall, these findings show that the Melanoma and ICI treatment dynamically alter microglial states, with evidence of both pro-inflammatory activation and disruption of homeostatic programs, underscoring the role of microglia as key mediators of neuroimmune alterations.

### ICI therapy disrupts Astrocytic homeostatic markers

We previously reported a modest decrease in the astrocytic marker GFAP following ICI treatment; however, the underlying transcriptomic dysregulation remained unclear. To better define astrocyte-specific transcriptional changes, astrocytes were identified using canonical gene markers and re-clustered with Seurat’s graph-based clustering, yielding multiple transcriptionally distinct subpopulations that were subsequently relabeled as A1–A18 for clarity (**Suppl. Fig. 3a**) “C:\Users\swami\Downloads\Supplemental Table Figures.docx”. Similar to the microglia, to assess relative distributions of these subpopulations under different experimental conditions, UMAP plots were split by the experimental groups (Control, Control + ICI, Melanoma, and Melanoma + ICI, **Fig. 4A)**. Similar to microglia, clustering of the astrocyte subset using the integrated astrocyte Seurat object at resolution 1.0 generated 18 astrocyte clusters, reflecting substantial transcriptional diversity. This resolution was selected because it captured meaningful astrocyte subpopulations while avoiding overfragmentation and maintaining cluster stability. The biological relevance of these clusters was further supported by heatmap analysis of astrocyte marker genes, which revealed cluster-specific expression profiles **(Suppl. Fig. S3b) “C:\Users\swami\Downloads\Supplemental Table Figures.docx“**. Pairwise Fisher’s exact tests revealed condition-specific shifts in astrocyte subclusters. Cluster A5 showed significant enrichment in melanoma-related comparisons (Control vs. Melanoma and Control vs. Melanoma+ ICI, P < 0.001; Control+ ICI vs. Melanoma and Control+ ICI vs. Melanoma+ ICI, P < 0.05). Cluster A8 displayed the strongest alterations, with highly significant differences in all melanoma-involving contrasts (P<0.001) and a modest but significant change in Control vs. Control+ ICI (P<0.001). Cluster A13 showed more limited changes, significant only between Control+ ICI vs. Melanoma and Control+ ICI vs. Melanoma+ ICI (P<0.05). Cluster A15 showed no significant differences (all P≥0.05). Overall, A5, A8, and A13 demonstrated condition-specific alterations, whereas A15 remained stable across datasets.

**Figure 4.**
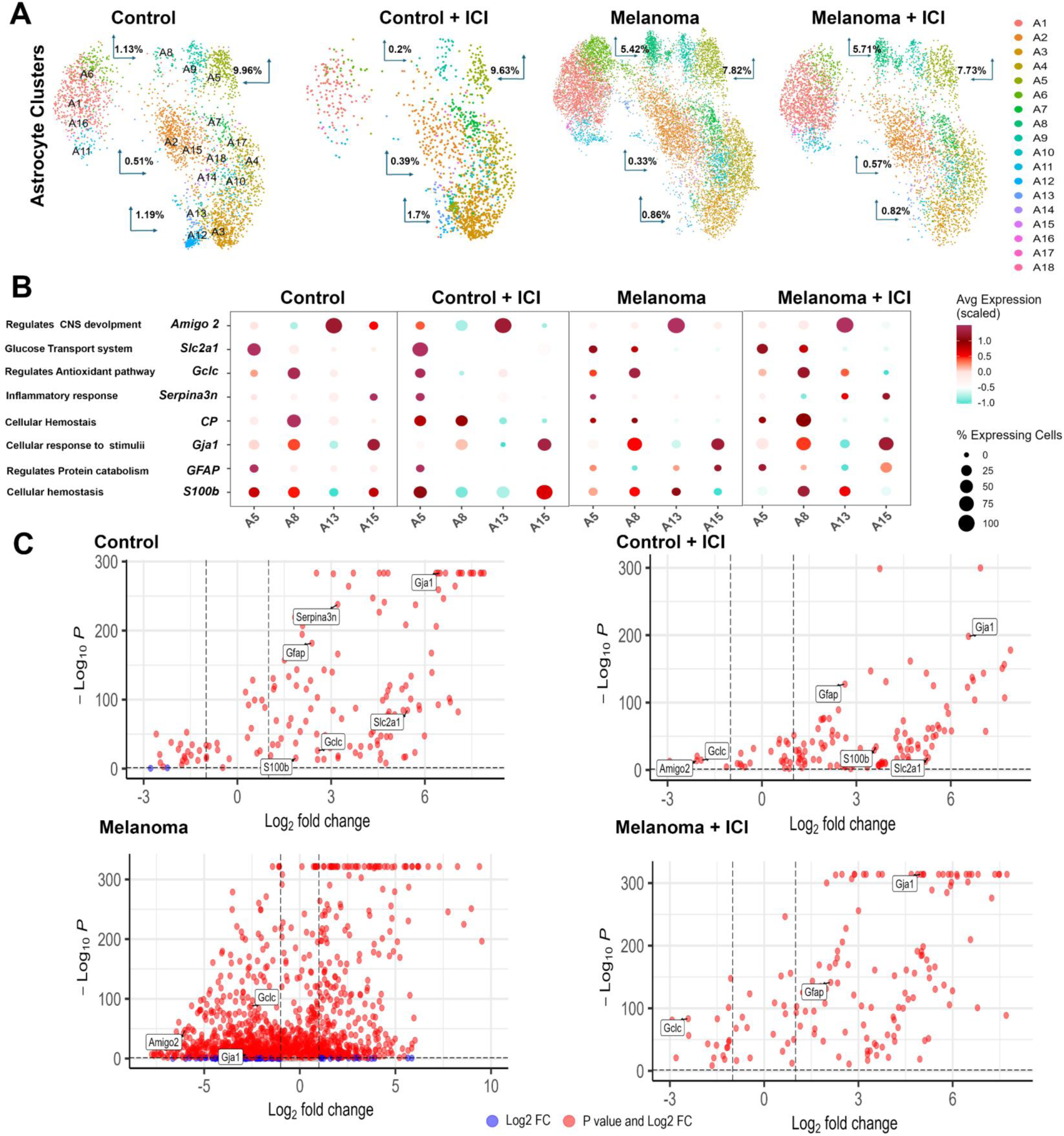
Gene expression analysis of the astrocyte cluster. **(A)** Split UMAPs display gene clusters across Control, Control +ICI, Melanoma, and Melanoma +ICI group datasets. **(B)** Bubble plots depicting the distribution and expression of genes within clusters. The size of the bubble represents the percentage of cells expressing the specific gene. The bubble color indicates the average expression level of that gene within the corresponding cluster. **(C)** Volcano plots depicting differential gene expression across the experimental group, with each point representing a gene. The X-axis corresponds to the log2 fold change (magnitude of expression difference between conditions), while the y-axis corresponds to the –log10 of the adjusted P-value (statistical significance after multiple testing correction, Benjamini-Hochberg FDR). Downregulated genes are further to the left, while upregulated genes are further to the right. Higher –log_10_*P* values indicated higher statistically significant gene expression. Significance thresholds used were: adjusted P<0.05 and log2 fold change > 1. Red: Strongly upregulated, high significance. Blue: Strongly downregulated, high significance Data are derived from N = 3 mice per group.

To quantify astrocyte cluster composition (A1-A18) across groups, we calculated the per-sample proportion of cells assigned to each cluster and summarized these distributions using 100% stacked bar plots **(Suppl. Table S3**) “C:\Users\swami\Downloads\Supplemental Table Figures.docx”. Further characterization of the functional identities of each cluster highlighted defining marker genes and their expression patterns across datasets **(Fig. 4B)**. Briefly, in the Control and Control + ICI groups, cluster A5 contained a higher proportion of cells (∼50%) with elevated expression of *Slc2a1* (GLUT1) compared to Melanoma and Melanoma + ICI groups, although its direct association with melanoma remains to be fully elucidated (Wang Y et al., 2023). Furthermore, the expression and proportion of cells expressing Gja1 increased across clusters A5, A8, A13, and A15 in the Melanoma and Melanoma + ICI groups relative to Control and Control + ICI, suggesting a role in mediating tumor–astrocyte or tumor–immune interactions (Hu W et al., 2020). Similarly, *GFAP* expression reached ∼75% across all clusters in the Melanoma + ICI group, consistent with astrogliosis following ICI treatment. In cluster A8, ∼75% of cells expressed *Cp* in the Control, Control + ICI, and Melanoma + ICI groups, suggesting restoration of neuroprotective functions after ICI therapy. Interestingly, cluster A15, which did not show significant differences across datasets, contained a high proportion (∼75%) of cells expressing S100b in the Control + ICI group, indicating stable expression of this neurotrophic marker across conditions.

To investigate gene-level insights complementary to the cluster-based observations, we next examined the expression of individual astrocyte markers across datasets **(Fig. 4C).** Differential expression analysis revealed that *Gja1*, a canonical astrocyte marker, was strongly upregulated in the Control (P<0.001, log2FC = 6.49), Control + ICI (P<0.001, log2FC = 6.56), and Melanoma + ICI (P<0.001, log2FC = 4.99) groups, but only moderately altered in the Melanoma group (P≥0.05).Similarly, *GFAP* (P<0.001, log2FC = 2.64 in Control + ICI) and *Slc2a1* (P<0.001, log2FC = 5.28 in Control + ICI), markers associated with reactive states and neuroprotective effects (Thieren L et al., 2025), were upregulated in Control + ICI and Melanoma + ICI, indicating astrogliosis responses in the context of ICI treatment and tumor presence. Conversely, Gclc (P < 0.05, log2FC = –2.42 in Melanoma + ICI), a gene linked to antioxidant pathways (Chen, Z et al., 2025), and Amigo2 (P < 0.001, log2FC = –6.01 in Melanoma + ICI), an anti-inflammatory marker (Luchena C et al., 2022), were downregulated in the Control + ICI and Melanoma + ICI groups, suggesting a dysregulation of astrocyte functional states under ICI conditions. Overall, these results highlight that astrocyte gene expression is modulated by ICI therapy, reflecting a balance between homeostatic maintenance and reactive activation.

### ICI therapy counteracts melanoma-associated oligodendrocyte dysregulation

To understand how ICI therapy affects myelination and contributes to the cognitive impairment reported previously, we examined the relative distributions of multiple transcriptionally distinct oligodendrocyte subpopulations, labeled O1–O17, using UMAP plots split by experimental group (Control, Control + ICI, Melanoma, and Melanoma + ICI, **Fig. 5A).** Similar to microglia and astrocytes, clustering of the oligodendrocyte subset using the integrated oligodendrocyte Seurat object at resolution 1.0 generated 17 oligodendrocyte clusters. The biological relevance of these clusters was further supported by heatmap analysis of oligodendrocyte marker genes, which revealed cluster-specific expression profiles (**Suppl. Fig. S4**b) “C:\Users\swami\Downloads\Supplemental Table Figures.docx”. Pairwise Fisher’s exact tests revealed condition-specific alterations in oligodendrocyte subclusters (**Suppl. Table S4**) “C:\Users\swami\Downloads\Supplemental Table Figures.docx”. Cluster O1 exhibited the strongest and most widespread shifts, with highly significant differences in all comparisons involving Control vs. Control+ ICI, Melanoma, and Melanoma +ICI (all adjusted P < 0.001). A significant difference was also observed between Melanoma and Melanoma+ICI (adjusted P < 0.05). Cluster O8 showed significant differences between Control vs. Melanoma and Control vs. Melanoma+ICI (adjusted P-values <0.001). Cluster O10 displayed significant changes in Control vs. Control+ ICI and Control+ ICI vs. Melanoma+ ICI (adjusted P<0.05). In contrast, Cluster O16 exhibited minimal variability, with only a modest difference observed between Control+ ICI and Melanoma+ ICI (adjusted P < 0.05), and no significant differences between Control and Melanoma. Overall, clusters O1, O8, and O10 demonstrated condition-specific shifts, whereas O16 remained largely stable across datasets.

**Figure 5.**
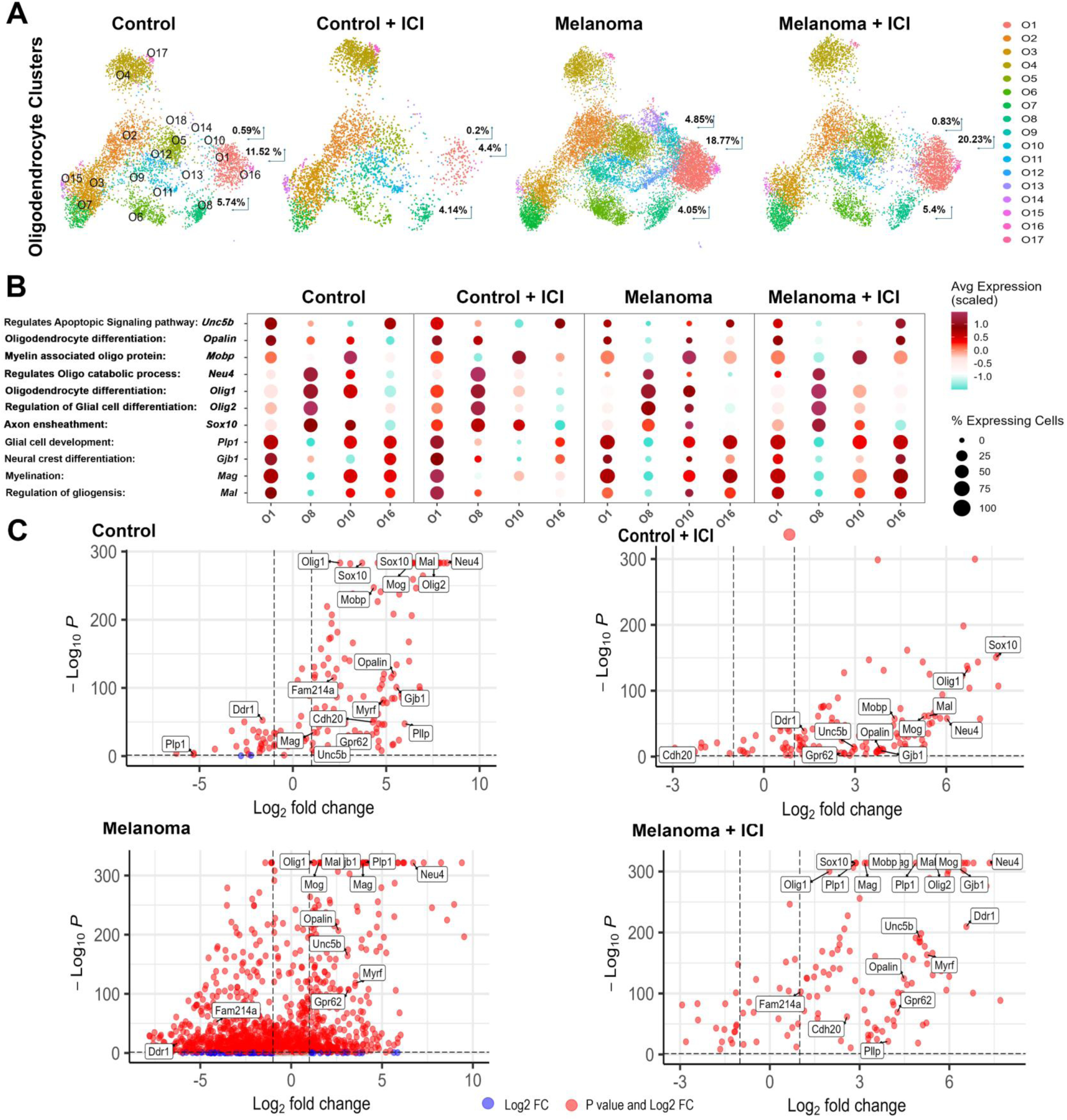
Gene expression analysis of the oligodendrocytes cluster. **(A)** Split UMAPs display gene clusters across Control, Control +ICI, Melanoma, and Melanoma +ICI group datasets. **(B)** Bubble plots depicting the distribution and expression of genes within clusters. The size of the bubble represents the percentage of cells expressing the specific gene. The bubble color indicates the average expression level of that gene within the corresponding cluster. **(C)** Volcano plots depicting differential gene expression across the experimental group, with each point representing a gene. The X-axis corresponds to the log2 fold change (magnitude of expression difference between conditions), while the y-axis corresponds to the –log10 of the adjusted P-value (statistical significance after multiple testing correction, Benjamini-Hochberg FDR). Downregulated genes are further to the left, while upregulated genes are further to the right. Higher –log_10_*P* values indicated higher statistically significant gene expression. Significance thresholds used were: adjusted P < 0.05 and log2 fold change > 1. Red: Strongly upregulated, high significance. Blue: Strongly downregulated, high significance Data are derived from N = 3 mice per group.

Furthermore, to characterize the composition of oligodendrocyte clusters (O1–O17) across conditions, we calculated the per-sample proportion of cells assigned to each cluster and visualized these distributions using 100% stacked bar plots (**Suppl. Fig. 4a** and **Suppl. Table S4**) “C:\Users\swami\Downloads\Supplemental Table Figures.docx”. Cluster O1 showed notable variation across datasets, driven in part by differential expression of *Unc5b*. Approximately 75% of cells in the Control and Control + ICI groups expressed Unc5b, compared with substantially lower proportions in the Melanoma and Melanoma + ICI groups **(Fig 5B**). Clusters O10 and O16 exhibited higher proportions of *Sox10-*expressing cells in the Control, Control+ ICI, and Melanoma+ ICI groups relative to Melanoma alone, indicating altered regulation of the oligodendrocyte-specific transcription factor *Sox1*0 in disease. Conversely, reduced expression of *Gjb1* within Cluster O1 in both Melanoma and Melanoma+ ICI groups suggests impaired CNS myelination and homeostasis under tumor and ICI-associated conditions (Papaneophytou et al., 2019).

To investigate gene-level changes complementing the cluster-based findings, we next examined the expression of individual oligodendrocyte markers across conditions **(Fig. 5C**). Differential expression analysis revealed consistent upregulation of core myelin-associated genes, including *Mag, Plp1, Mog, Mal, and Opalin*, in the Control, Control + ICI, and Melanoma + ICI groups (all P<0.001), indicating preservation of mature oligodendrocyte identity across these conditions. In contrast, the Melanoma group showed a more restricted expression profile, with *Ddr1 and Fam214a* significantly downregulated (both P<0.05). The Control + ICI group exhibited downregulation of only *Cdh20*gene. This pattern indicates that myelin-related transcriptional programs remain intact under ICI therapy. Notably, several genes (*Sox10, Myrf, and Unc5b*) were also robustly upregulated in Melanoma + ICI, suggesting that ICI may partially reverse or compensate for melanoma-induced oligodendrocyte dysfunction. Overall, these results demonstrate that oligodendrocyte gene expression is differentially affected by melanoma and by ICI treatment

### ICI therapy stimulates T Cell activation and NF-κB-mediated inflammatory responses

To assess the relative distributions of the transcriptionally distinct T cell subpopulations, UMAP plots were split by experimental groups (Control, Control + ICI, Melanoma, and Melanoma + ICI; **Fig. 6A**). Pairwise Fisher’s exact tests revealed condition-specific alterations across several T cell clusters (**Suppl. Table S5**). Cluster T7 exhibited highly significant differences across the datasets. Comparisons of Control vs. Control + ICI, Melanoma, and Melanoma + ICI all showed strong shifts (P<0.001). A significant alteration was also observed between Melanoma vs. Melanoma + ICI (P<0.001), indicating that T7 abundance is influenced by both melanoma and ICI treatment. Additionally, Cluster T5 showed a moderate shift across the datasets, with significant differences between Control vs. Melanoma (adjusted P < 0.05) and Melanoma vs. Melanoma + ICI (adjusted P < 0.01). Cluster T14 showed significant differences between Control vs. Melanoma and Control vs. Melanoma + ICI (P<0.05). Conversely, Cluster T11 showed no significant differences across any of the datasets, suggesting no major transcriptional changes between treated and untreated groups

**Figure 6.**
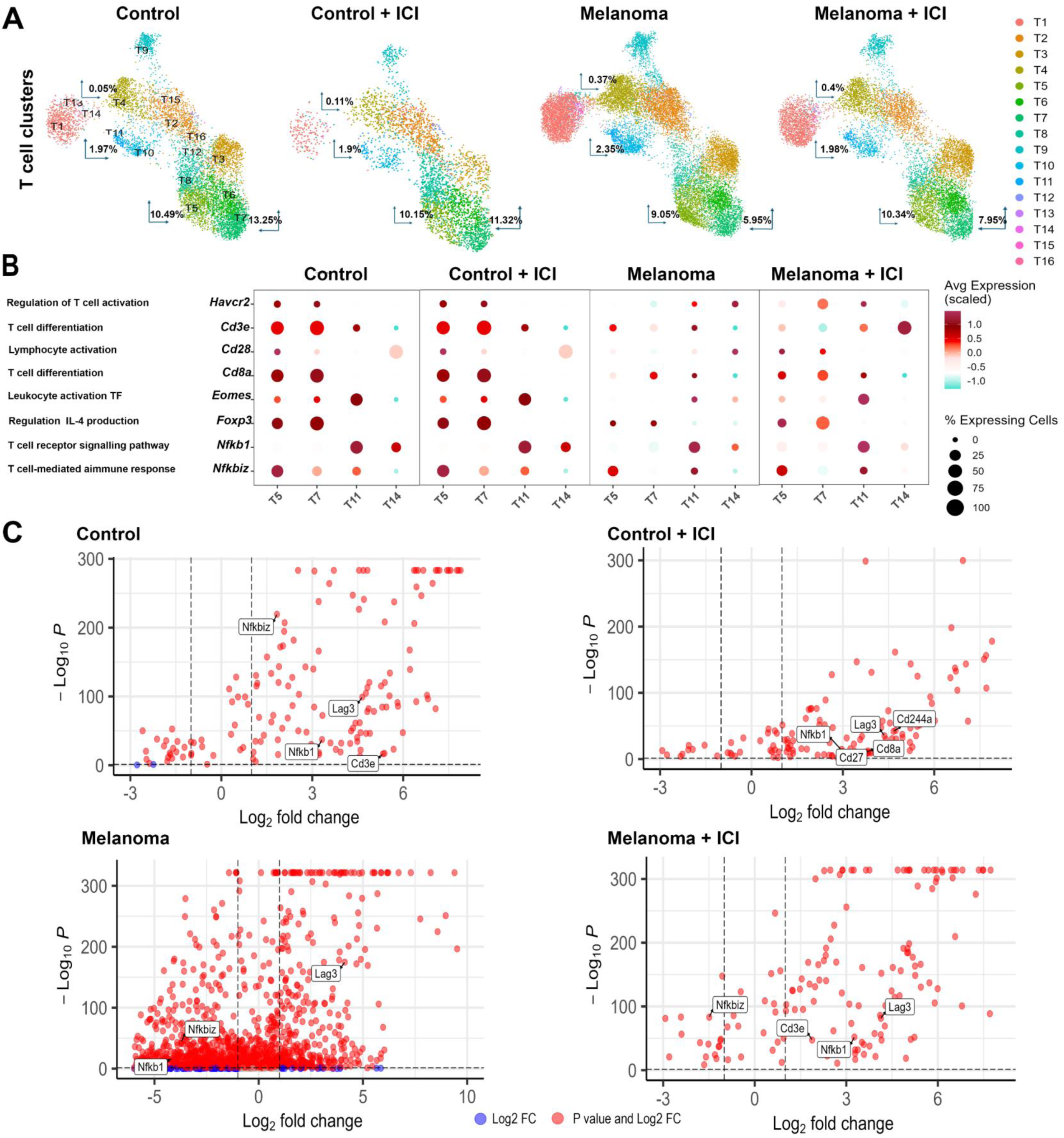
Gene expression analysis of the T cell cluster. **(A)** Split UMAPs display gene clusters across Control, Control +ICI, Melanoma, and Melanoma +ICI group datasets. **(B)** Bubble plots depicting the distribution and expression of genes within clusters. The size of the bubble represents the percentage of cells expressing the specific gene. The bubble color indicates the average expression level of that gene within the corresponding cluster. **(C)** Volcano plots depicting differential gene expression across the experimental group, with each point representing a gene. The X-axis corresponds to the log2 fold change (magnitude of expression difference between conditions), while the y-axis corresponds to the –log10 of the adjusted P-value (statistical significance after multiple testing correction, Benjamini-Hochberg FDR). Downregulated genes are further to the left, while upregulated genes are further to the right. Higher –log_10_*P* values indicated higher statistically significant gene expression. Significance thresholds used were: adjusted P < 0.05 and log2 fold change > 1. Red: Strongly upregulated, high significance. Blue: Strongly downregulated, high significance Data are derived from N = 3 mice per group.

Additionally, the proportion of cells per sample was calculated for each cluster, and their distribution was visualized using 100% stacked bar plots (**Suppl. Fig. 5a** and **Suppl. Table S5**) “C:\Users\swami\Downloads\Supplemental Table Figures.docx”. The biological relevance of these clusters was further supported by heatmap analysis of T cell marker genes, which revealed cluster-specific expression profiles (**Suppl. Fig. 5b) “C:\Users\swami\Downloads\Supplemental Table Figures.docx“** In **Fig. 6B**, Cluster T14 showed nearly 100% of cells expressing Havcr2 in the Control + ICI group compared to the Melanoma and Melanoma + ICI groups, suggesting enhanced T cell activation following ICI therapy (Alamir H. et al., 2025). Similarly, *Cd3e* (Clusters T5, T7), *Cd28* (Cluster T14), and *Cd8a* (Clusters T5, T7) were expressed by approximately 75% of cells in the Control and Control + ICI groups, whereas their expression was markedly reduced in the Melanoma and Melanoma + ICI groups. In contrast, the *Nfkbiz* gene showed substantial variability across all clusters (T5, T7, T11, and T14) and datasets.

To investigate gene-level changes complementing the cluster-based findings, we next examined the expression of key T cell–associated genes across all four conditions (**Fig. 6C**). Differential expression analysis revealed robust upregulation of canonical markers associated with T-cell signaling and activation, including *Cd3e, Lag3, Nfkb1, and Nfkbiz* in the Control group, indicating a stable baseline activation and homeostatic regulatory state in resting T cells. In contrast, the Melanoma group displayed a markedly restricted transcriptional profile, characterized by strong downregulation of *Nfkb1* (P<0.05), a pattern consistent with tumor-driven T cell exhaustion and impaired NF-κB–mediated inflammatory signaling (Caratelli S. et al., 2025). Notably, ICI treatment in the Melanoma + ICI group, partially restored T cell activation programs. Genes such as *Nfkb1 and Cd3e* were also expressed at higher levels, while *Lag3* remained highly upregulated, indicating persistent but modulated T cell activation post ICI therapy. The modest downregulation of *Nfkbiz* in this group is consistent with altered NF-κB regulatory feedback associated with inflammatory activation post ICI therapy (Feng Y et al., 2023)

### Immunofluorescence analyses of postmortem brains from patients treated with ICI

To assess whether non-neuronal gene programs identified in our mouse MERFISH dataset translate to human neurotoxicity, we selected a panel of cell–type–specific markers shortlisted from MERFISH profiling of mouse brains across experimental groups (Control, Control+ICI, Melanoma, and Melanoma+ICI) and evaluated their expression in frontal cortex from postmortem human brain tissue. We compared postmortem tissue from patients receiving ICI treatment (anti-PD-1, and Combi-ICI) for different malignancies to a noncancer, non-treated control cohort. Using dual immunofluorescence staining, we measured the immunoreactivity of cell type–specific markers, including microglial activation (*CD16/FCGR3 with TREM2*), astrogliosis (*GFAP with SERPINA3*), and oligodendrocytes (*SOX10 with MBP*) in the frontal cortex of postmortem patients **(Fig. 7)**. Quantification of single and co-localized immunoreactivity was performed using an *in silico* 3D algorithm-based volumetric approach as described previously (Markarian et al., 2021). Patients treated with anti-PD-1 therapy (ICI patients) showed a significant increase in dual-labeling of *CD16/FCGR3 and GFAP/SERPINA3* immunoreactivity volumes compared with non-cancer control patient brains (P<0.0001). In contrast, *MBP* immunoreactivity was reduced by approximately 20% following anti-PD-1 treatment compared with non-cancer controls (P < 0.05). *GFAP/SERPINA3* co-localization was elevated by 1.5 times in non-cancer control patients compared with anti–PD-1-treated patients. These findings corroborate the transcriptomic data demonstrating upregulation of *CD16/FCGR3, GFAP, and SOX10,* along with reduced expression of myelin markers in melanoma + ICI-treated mice. Collectively, these data indicate a pronounced dysregulation of microglial, astrocytic, and oligodendrocyte markers following ICI therapy, as supported by converging evidence from human patient brains treated with ICI.

**Figure 7.**
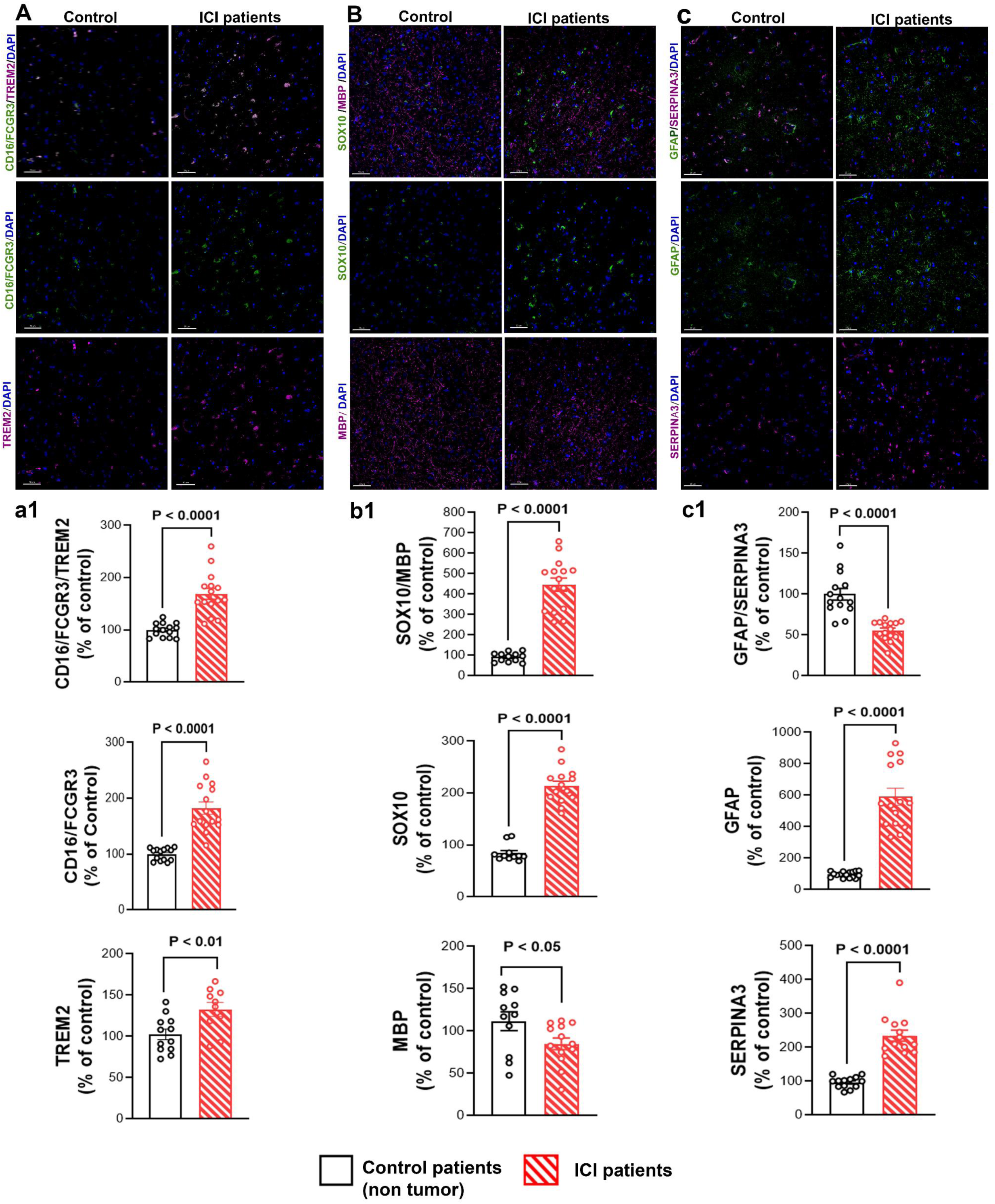
Immunofluorescence analysis of human postmortem brains from ICI-treated patients. **(A–C)** Dual immunofluorescence staining for CD16/FCGR3 and TREM2 (microglia), GFAP and SERPINA3 (astrocytes), and SOX10 and MBP (oligodendrocytes and myelin) in frontal cortices from N=3 patients receiving ICI treatment for different malignancies compared with N=3 non-cancer, untreated control cohort. Scale bar = 50µm **(a1–c1)** Volumetric quantification of immunoreactivity from 3-4 region of interest (ROI) per brain section (N=4 sections per brain) corresponding to **(A-C)** shows that ICI-treated patient cortices had increased immunoreactivity for microglial-, astrocyte-, and oligodendrocyte-specific gene markers compared with non-cancer controls. Data are represented as Mean ± SEM, each data point represents one region of interest per brain section. N = 4 sections from each ICI patients and control non tumor brains. P values were derived from unpaired, two tailed *t* tests.

**Figure 8.**
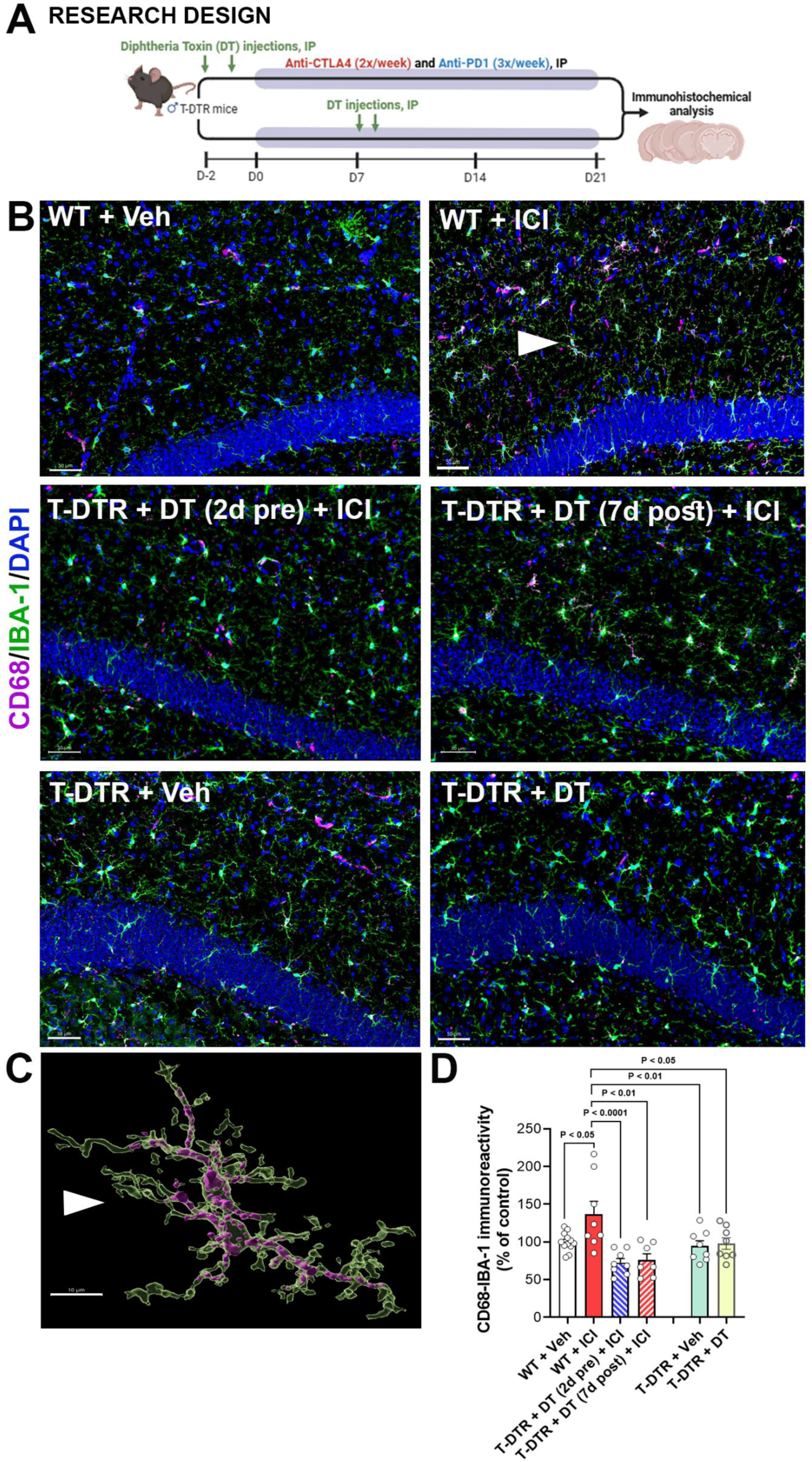
Depletion of T cells prevents ICI treatment-induced microglial activation. **(A)** Experimental Design: Transgenic T-DTR mice received diphtheria toxin A (DT) either 2 days before or 7 days after ICI treatment (anti-CTLA-4, and anti-PD-1). ICI treatments were given for 2 weeks (see Materials & Methods for details). WT (C57Bl/6) mice receiving either ITC or ICI treatment were used for comparison. Brains were collected at 72 hours after completion of ICI treatments. **(B)** Representative dual-immunofluorescence confocal z stacks depicting microglial activation (CD68, magenta, IBA1, green, and DAPI nuclear counterstaining) from T-DTR and WT mouse hippocampus. Scale bar = 50 µm **(C)** A 3D surface rendering of CD68-IBA1^+^ activated microglia from the WT + ICI group (white arrows). Data are presented as Mean ± SEM, N=6-8 mice/group. P values were derived from ANOVA and Dunn’s multiple comparisons test. **(D)** Volumetric quantification of CD68 and IBA1 immunoreactivity showing ICI treatment elevated microglial activation in WT + ICI groups compared to WT + Vehicle. T cell depletion (T-DTR mice) did not lead to microglial activation.

### Conditional T cell depletion reduces microglial activation following ICI treatment

Based on previously proposed mechanisms (Wei et al., 2018, Buchbinder EI, et al., 2016) the synergistic blockade of CTLA-4 and PD-1 in ICI therapy may (i) unleash T cells and trigger a cascade of neuroinflammation, (ii) disrupt neuroimmune homeostasis, thereby promoting microglial activation. To further test this neuroimmune crosstalk, we examined how T cells influence microglial status, using a transgenic mouse model harboring an inducible diphtheria toxin receptor (iDTR) transgene (C57BL/6-Gt(ROSA)26Sortm1(HBEGF)Awai/J; B6-iDTR). The cross of B6-iDTR mice with CD4-Cre transgenic mice (or T-DTR) results in offspring to conditionally ablate CD4⁺ and CD8⁺ T lymphocytes using diphtheria toxin A (DT) treatment. To achieve robust T cell depletion, T-DTR mice were treated with DT (100 ng, once daily for 2 days) at 2 days before or 7 days after initiation of ICI treatment (**Fig. 8A**). ICI treatments were given for two weeks (see **Materials and Methods** for details). Dual immunofluorescence staining for microglial activation (CD68-IBA1, **Fig. 8A**,) was conducted in brains collected at 72 hours after either ICI or DT treatments. CD68 is a lysosomal membrane protein and an indicator of phagocytosis, and IBA1 is a pan-microglial marker. WT mice (C57Bl/6) receiving vehicle or ICI treatment were used for comparison. ICI treatment to the WT mice showed a significant elevation in the CD68⁺ and IBA1⁺ activated microglia compared with vehicle-treated WT mice (P<0.05, **Fig. 8B-D, Suppl. Fig. 5**) “C:\Users\swami\Downloads\Supplemental Table Figures.docx”. DT- and vehicle-treated T-DTR mice (without T cell depletion) did not show any changes in the CD68-IBA1 immunoreactivity. In contrast, T cell depletion in T-DTR mice at 2 days before or 7 days after ICI treatment did not elevate microglial activation, and the CD68-IBA1 dual-immunoreactivity was comparable to that of WT mice receiving vehicle. These data show that T cells are required for ICI-related microglial dysregulation and support the hypothesis that ICI-unleashed T cells are the key drivers of microglial activation during ICI therapy.

## DISCUSSION

Our study reports a comprehensive spatial transcriptomic profile of cell–type–specific markers and their associated regulatory networks following ICI therapy. Following ICI treatment, we identified coordinated cell–type–specific changes associated with neuroinflammation, including inflammatory activation of microglia, reactive astrogliosis, reduced myelin integrity in oligodendrocytes, and an NF-κB–associated signature in T cells. Translational validation using postmortem brain tissue from human patients treated with anti-PD-1 therapy corroborates transcriptomic findings. Mechanistically, converging evidence from T cell-depleting experiments demonstrates that ICI-driven microglial activation depends on T cells. Collectively, our findings not only reveal cellular and molecular networks that underlie ICI-mediated cognitive decline but also establish ICI-unleashed T cells as the key drivers of microglial dysregulation during ICI therapy.

### Hippocampal RNA-seq reveals coordinated T cell, microglial, and astrocyte activation signatures

Bulk RNA sequencing of hippocampus revealed robust upregulation of microglial-, astrocytic-, and T cell–associated neuroinflammatory markers in both Control + ICI and Melanoma + ICI groups, indicating a sustained inflammatory response following ICI therapy. Among the significantly upregulated genes common to both ICI-treated groups, *Adora2a* (adenosine A2A receptor) emerged as a key candidate linked to neuromodulatory signaling. Two of the four adenosine receptors, A1R and A2AR, are highly expressed throughout the brain and mediate adenosine’s, neuro-modulatory effects (Lemes Dos Santos Sanna P et al., 2024). Notably, A2AR is expressed in astrocytic processes, microglia, and activation of A2AR has been associated with neurodegenerative processes; in the hippocampus, it enhances neuronal excitability and can promote neuronal injury and cell death (Ikram et al., 2020; Stockwell et al., 2017). Empirical studies further demonstrate that A2AR density and immunoreactivity increase in hippocampal nerve terminals during aging in rodents (Rebola et al., 2003) and in humans (Temido-Ferreira et al., 2020). Importantly, this overexpression has been linked to neuroinflammation and memory impairment (Launay A et al., 2020). Consistent with these findings in our dataset, *Adora2a* expression was elevated in Melanoma + ICI, supporting a notion that the adenosine–A2A axis has a potential role in astrocytic dysfunction and neuroinflammation following ICI treatment.

Furthermore, the top differentially expressed genes (DEGs; P < 0.05) across datasets were identified, including microglia- and synapse-related markers that were upregulated in both the Melanoma and Melanoma + ICI groups. Prior studies have shown that impaired MHC class II expression can lead to profound immune perturbations associated with CD4⁺ T cells (Jurewicz MM et. Al., 2019). *H2-Eb1, H2-Ab1, and H2-A*a encode the β and α chains of murine MHC class II (I-A/I-E) and are canonical markers of MHC II–high activated microglia/macrophages in the CNS (Majumder P et al., 2021). Upregulation of these genes in microglia or tumor-associated macrophages (TAMs) is typically interpreted as enhanced antigen-presenting capacity, which promotes CD4⁺ T-cell priming and support anti-PD-1/anti-CTLA-4 responses in brain metastases (Logunova N et al., 2023). Consistent with this, other studies on genomic profiling of 206 patients with advanced melanoma treated with immune checkpoint blockade (anti-PD-1, anti-CTLA-4, or combination therapy) showed higher expression of MHC II-associated genes in responders, alongside enrichment of immune-related pathways, including inflammatory response and IL-6/JAK/STAT3 signaling (Liu D et al., 2019). Together, these findings suggest persistent inflammatory signatures post-ICI therapy.

Similarly, the upregulation of *Darp-32* in the Melanoma + ICI group may imply a role in ICI-induced cognitive impairment, consistent with findings from previous studies. Dopamine- and cAMP-regulated phosphoprotein of 32 kDa (*Darp-32)* plays a central role in dopaminergic signaling. Phosphorylation of DARPP-32 in response to L-DOPA regulates its dual function as either an inhibitor or facilitator of protein phosphatase-1 (PP1), thereby modulating neuronal excitability and synaptic plasticity (Girault JA et al., 2021). L-DOPA–induced activation of *Darp-32* has been strongly associated with L-DOPA–induced dyskinesia (LID), particularly in medium spiny neurons (MSNs), highlighting the critical role of the cAMP/ *Darp-32* signaling cascade in neuronal dysfunction (Santini E et al., 2012). In our study, *Darp-32* was significantly upregulated in ICI-treated groups, suggesting a potential link between altered dopaminergic signaling and cognitive impairment following ICI therapy. However, further validation of additional dopaminergic pathway markers within the transcriptomic dataset is necessary to better elucidate the mechanistic relationship between *Darp-32* upregulation and ICI-induced neurocognitive alterations.

Evaluation of transcriptomic signatures across datasets further identified several strongly upregulated genes, including *Tmem252, Lcn2, Apold1, Rarb, Ptprv, Adora2a, and Cd4,* in the Melanoma + ICI group. Consistent with these findings, violin plot analyses demonstrated that *Ptprv, Adora2a, and Cd4* were significantly elevated in both the Melanoma + ICI and Melanoma groups. Higher *CD4* expression was reported in responders within the ipilimumab-treated subgroup (Liu et al., 2019), supporting a role for immune activation in the context of ICI therapy. In our experimental dataset, ICI treatment significantly increased *Ptprv* expression in both Melanoma and Melanoma + ICI groups. This dysregulation aligns with prior reports showing that *Ptprv* transcription is markedly and preferentially induced in cells undergoing p53-dependent cell cycle arrest (Doumont et al., 2005). The p53 transcriptional activity has also been shown to modulate microglial pro-inflammatory functions while suppressing anti-inflammatory and tissue repair responses (Aloi et al., 2015). Collectively, these findings suggest that the *Ptprv*-p53 axis potentially contributes to microglial activation and neuroinflammation after ICI therapy.

### Cross-species validation of spatial gene signatures reveals ICI-linked neuroinflammation across glia and T Cells

Disturbance in CNS homeostasis (e.g., neuroinflammation) triggers microglial activation with morphological changes and increased inflammatory signaling, while astrocytes can adopt a neurotoxic reactive state (Garland EF et al., 2022). Moreover, oligodendrocytes also actively participate by interacting with microglia, astrocytes, and infiltrating peripheral immune cells, altering their signaling programs and releasing pro-inflammatory cytokines such as TNF-α, IL-1β, and IL-6 (Pasquini JM et al., 2025). Together, these observations emphasize that neuroinflammation is network-driven, arising from coordinated crosstalk among microglia, astrocytes, and oligodendrocytes. However, how these interactions among non-neuronal cells and inflammatory immune cells are altered by ICI therapy was largely undefined. Our integrated MERFISH analysis provides new insights into markers across oligodendrocytes, astrocytes, T cells, and microglia, emphasizing the need to (i) delineate multi-cellular interactions, (ii) for a better understanding of ICI-associated neuroinflammatory and neurodegenerative sequelae, and (iii) to guide safer therapeutic strategies. Our microglial sub-clustering indicates that ICI therapy drives microglia toward activated and immune-regulatory states consistent with previous reports (Vinnakota JM et al., 2024), and prior findings (Ifejeokwu et al.,2025). In the Control + ICI group, microglial clusters M4 and M11 showed elevated *Aif1* and *Fcgr3*, supporting a shift toward an activated phenotype. This pattern aligns with published work describing *Aif1* as a dynamically regulated, injury-associated marker that can act as an early sensor of acute and delayed neuronal damage (Schwab JM et al., 2020). Similarly, increased *Fcgr3/Cd16* is consistent with studies linking *Fcgr3a* to immune signaling and angiogenesis-related pathways, and with higher *Fcgr3a* expression being associated with altered responses to anti-tumor therapies (Li L et al., 2022).

Furthermore, in the Melanoma + ICI group, the expanded M8 microglial cluster exhibited increased *Cd300lf*, a regulator of microglial responses to tissue damage. Notably, this finding contrasts with prior work reporting a neuroprotective role for *Cd300lf*, potentially via the STING signaling pathway, in models of brain injury (Lu Z et al., 2022). Collectively, these results indicate that ICI elicits a heterogeneous microglial response, with concurrent induction of inflammatory and immune-modulatory pathways rather than a uniform activation state. Importantly, validation in postmortem human brains showed a higher Fcgr3/CD16 signal in anti-PD-1–treated patients than in noncancer controls, further supporting an inflammatory microglial signature following ICI therapy.

Complementing these observations, our gene-level analysis showed that *Gpr84* (a microglia-associated pro-inflammatory GPCR) was significantly upregulated in both the Melanoma and Melanoma + ICI groups. GPCRs are key mediators of neuron-glia communication under pathological stress (Wei L et al., 2017), and mechanistically, increased *Gpr84* expression in microglia has been linked to NLRP3 inflammasome activation, potentially through inhibition of the cAMP/PKA signaling pathway (Jiang K et al., 2025).Together, these findings indicate that ICI treatment drives microglia toward inflammatory/immune-regulatory states (*Aif1/Fcgr3/Cd300lf*) and induces sustained *Gpr84* signaling, suggesting an elevated neuroinflammatory state following ICI therapy.

Astrocyte clustering revealed a pronounced shift toward a reactive astrocyte phenotype in tumor-bearing and ICI-treated conditions. Across astrocyte clusters, a large proportion of cells expressed *Gja1 and Gfap*, with the strongest *Gfap*-associated reactivity observed in the Melanoma and Melanoma + ICI groups. *Gja1*, also known as connexin 43 (Cx43), is a member of the connexin family of gap junction proteins, and chronic elevation of *Gja1/Cx43* has been associated with neurodegenerative disease progression and cognitive impairment, with sustained upregulation potentially exerting detrimental effects on neurons (Kajiwara Y et al., 2018). In parallel, clinical observations further support our findings: ICI-treated patients have been reported to harbor anti-GFAP autoantibodies in cerebrospinal fluid (CSF), implicating astrocyte-directed immune responses in immune-related adverse events (Buckley MW et al., 2025; Kapadia RK et al., 2020). Consistent with these reports, our gene-level analyses also showed *Gfap* upregulation in ICI-treated experimental groups, suggesting that ICI-associated astrocyte changes are driven by reactive-state programs.

Importantly, when translated into a human context, anti-PD-1-treated patient brains showed dysregulated astrocyte markers compared with noncancer controls, including significantly increased *Gfap* and *Serpina3* immunoreactivity. Together, these cross-species data support the conclusion that ICI is associated with heightened *Gfap*-linked astrocyte reactivity,—indicating that ICI-related astrocyte responses are dominated by a reactive inflammatory state rather than restoration of astrocyte homeostasis.

Oligodendrocyte sub-clustering indicated that ICI therapy alone exerts a comparatively modest effect on core oligodendrocyte and myelin programs, whereas melanoma is associated with more pronounced dysregulation. In particular, *Unc5b* expression was higher in the Control and Control + ICI groups but reduced in Melanoma and Melanoma + ICI. This pattern is consistent with prior reports showing that *Unc5b* is enriched in mature oligodendrocytes and is required for myelin sheath stability (Faria et al., 2025), supporting the interpretation that tumor-bearing conditions are linked to compromised oligodendrocyte homeostasis and myelin maintenance. We also observed cluster-specific variation in *Sox10* expression across oligodendrocyte clusters (O1, O8, O10, and O16) in the dataset, with O8 exhibiting the highest *Sox10* expression. The elevated *Sox10* in the ICI group is consistent with prior reports linking *Sox10* to immune checkpoint regulation: Yokoyama and colleagues (2021) reported that *Sox10* suppresses PD-L1 expression via the IRF4–IRF1 axis, whereas Sasaki et al. (2021) reported increased PD-L1 expression following *Sox10* overexpression. Further studies are warranted to clarify the relationship between *Sox10* and immune checkpoint signaling. In addition, we detected dysregulation of *Ddr1* across conditions: compared with Control and Control + ICI, the Melanoma group showed significant downregulation of *Ddr1*. Given DDR1’s established roles in extracellular matrix interactions, cell adhesion, and myelin remodeling (Zhang J et al., 2025), reduced *Ddr1* further supports melanoma burden-associated impairment of oligodendrocyte structural and functional stability. Consistent with the notion that ICI alone has limited effects on baseline myelin programs, core myelin genes (*Mag, Plp1, Mog, Mal,* and *Opalin*) remained relatively stable between Control and Control + ICI. Translationally, anti-PD-1-treated patient brains also showed elevated SOX10 immunoreactivity, which differs from patterns observed in the mouse oligodendrocyte compartment and may reflect differences in disease stage, region, or cellular composition across species. Importantly, the reduction of MBP immunoreactivity in human ICI patients brains aligns with the myelin-instability signature suggested by reduced *Unc5b* in the mouse dataset. Previously, we showed reduced MBP immunoreactivity in the brains of Melanoma + ICI mice (Ifejeokwu OV et al., 2025). Collectively, these findings support myelin dysregulation in ICI-associated neurotoxicity.

Our spatial transcriptomic profiling of T cell markers revealed a distinct shift in T cell programs in response to melanoma and ICI therapy. In the Melanoma and Melanoma + ICI groups, we observed reduced expression of *Cd3e, Cd28, and Cd8a*, key genes involved in T cell development, activation, and differentiation. Functionally, *Cd3* subunits mediate signal transduction required for T cell activation, while *Cd28* regulates co-stimulatory signaling pathways that support T cell activation and survival. Prior studies have shown that *Cd28* downregulation is associated with severe T cell senescence and may contribute to resistance to immune surveillance and immunotherapy, leading to poor clinical outcomes (Humblin E et al., 2023). Moreover, sustained CD28 co-stimulation is required for the maintenance and expansion of TCF1⁺PD1⁺CD8⁺ T cells (Huang Y et al., 2023). Together, these findings suggest that the reduced expression of T-cell activation markers following ICI therapy reflects a shift in T-cell transcriptional programs within the neuroinflammatory brain microenvironment, potentially indicating altered activation states rather than classical cytotoxic T-cell responses

In contrast, the Melanoma + ICI group showed a higher proportion of cells expressing *Nfkbiz*, suggesting an ICI-linked, T cell-associated inflammatory response (Feng Y et al., 2023). Supporting this, our gene-level analysis also indicated increased expression of *Nfkb1* in the Melanoma + ICI condition. NF-κB (including the gene variants *Nfkb1 and Nfkbiz*) is a transcription factor that plays a central role in neuroinflammation. Increased NF-κB activity, often reflected by elevated pro-inflammatory cytokine expression, can contribute to neuronal degeneration (Momtazmanesh et al., 2020; Shastri et al., 2013). Conversely, inhibition of NF-κB has been shown to reduce neural loss and improve cognitive function by attenuating pro-inflammatory cytokine signaling (Srinivasan and Lahiri, 2015). Additionally, NF-κB signaling are critical in determining T cell differentiation, responses and fate (Daniels MA et al., 2025These prior findings corroborate our spatial transcriptomic analysis of T-cell markers, which reveals enhanced NF-κB–associated inflammatory signaling following ICI therapy. However, further validation in ICI-treated samples using T-cell markers such as CD4 and CD8 will strengthen the cross-species relevance of our transcriptomic data. Together, our spatial profiling indicates that post-ICI T cells display an NF-κB–associated inflammatory activation signature (increased *Nfkbiz/Nfkb1*) despite reduced expression of canonical activation/cytotoxic markers (decreased *Cd3e/Cd28/Cd8a*).

### ICI-driven microglial activation requires T cells

Immune checkpoint inhibitors (ICIs) enhance antitumor immunity by blocking the inhibitory pathways of CTLA-4 and PD-1, thereby unleashing peripheral and tissue-resident T-cell responses. Although this mechanism is therapeutically beneficial against cancer, it can also disrupt neuroimmune homeostasis in the CNS. Based on this mechanism, we conceptualize that combined ICI unleashes T cells while concurrently disrupting the PD1–PDL1 axis in the brain, leading to microglial activation and potentiation of local neurodegenerative inflammatory loops. In our study, transcriptomic profiling supported this concept by revealing coordinated alterations in both T cells and microglia following ICI exposure. Previous work showed that T-cell depletion in TE4 mice reduced microglial MHC class II and CD11c expression, decreased microglial immunoreactivity and brain atrophy, and improved short-term memory (Chen X. et al., 2023). However, whether microglial reactivity during ICI therapy is driven by T cells remains unexplored. In our study, conditional T-cell depletion in T-DTR mice, performed either 2 days before or 7 days after ICI treatment, prevented ICI-induced microglial activation, as CD68/IBA1 dual immunoreactivity remained low and comparable to WT vehicle-treated controls. In contrast, WT mice with an intact T-cell compartment showed marked microglial activation following ICI treatment. These findings suggest that ICI first unleashes T cells, which subsequently drive microglial activation. Consistent with this interpretation, T-DTR mice treated with DT showed no significant microglial activation compared with ICI-treated WT mice. Together, these results demonstrate that conditional T-cell depletion prevents ICI-mediated microglial activation.

These findings are consistent with previous reports showing that CAR19 T cell–treated mice exhibited increased numbers of IBA1⁺ microglia in the cortex compared with mice treated with natural killer T cells, indicating that microglial activation is induced in the presence of T cells (Vinnakota J.M. et al., 2024). Moreover, Chen and colleagues (Chen X. et al., 2023) demonstrated that blocking T-cell immunoregulation through repeated anti-PD-1 administration in TE4 mice altered immune homeostasis. Collectively, these studies indicate that T-cell activation drives microglial reactivity and disrupts neuroimmune homeostasis in the CNS. Our findings therefore support a coherent model in which ICI treatment unleashes T-cell activation and drives secondary microglial reactivity in the brain. The altered expression of microglial homeostatic markers together with elevated T-cell activation markers in our transcriptomic analyses supports the presence of an active T cell–microglia inflammatory axis. Building on this observation, our T-DTR depletion experiments provided functional validation of this axis by demonstrating that removal of T cells attenuated microglial activation following ICI treatment. Collectively, these results identify T cells as key upstream drivers of microglial reactivity and suggest that limiting T cell–mediated neuroinflammation may represent a protective strategy against ICI-associated neurodegenerative changes.

A key limitation of our study is that we focused primarily on changes in gene expression and cellular composition in the hippocampus related to cognitive impairment following ICI therapy. However, future studies investigating interactions between non-neuronal cells and inflammatory immune cells in other brain regions, particularly the prefrontal cortex, choroid plexus, and meninges, are warranted to better understand how these crosstalk networks contribute to neuroinflammation and cognitive dysfunction after ICI treatment. In addition, spatial transcriptomic profiling of brain tissue from patients receiving ICI therapy is needed, as it remains unclear to what extent the cell–type–specific markers identified in our experimental mouse model are conserved in the human brain. Potential molecular pathways (e.g., IFNγ and TNFα) of the T cell-microglial crosstalk also need further investigation, and another open question is to what extent T cell depletion can rescue ICI-related cognitive dysfunction.

In summary, our work defines cellular and molecular networks that drive neuroinflammation following ICI therapy. Translational validation of these transcriptomic signatures—particularly increased expression of key neuroinflammatory markers (Cd16/Fcgr3, Gfap, and Sox10) together with reduced expression of myelin stability markers—supports coordinated crosstalk among microglia, astrocytes, and oligodendrocytes after ICI therapy. In addition, our findings clearly demonstrate that T cells are essential players in ICI-driven microglial activation. Our study not only establishes a robust mechanistic framework for how ICI therapy alters neuroinflammatory pathways but also provides opportunities to develop strategies to decouple microglial activation from the anti-tumor effects of ICI therapy.

## Supporting information

"C:UsersswamiDownloadsSupplemental Table Figures.docx"

## ACKNOWLEDGEMENTS

We thank Prof. T. Z. Baram and M. Tetzlaff for their support of the MERFISH experiments. We thank A. H. Do, A. Sundar, K. M. C. Nguyen, O. V. Ifejeokwu, R. P. Krattli, S. Madan, S. M. El-Khatib, and T. Nguyen for their logistical and technical assistance. We thank Dr. F. Marangoni for kindly providing the syngeneic murine melanoma cell line and guidance.

## DECLARATIONS

- Ethics approval: All animal experiments were approved by the UCI Institutional Animal Care and Use Committee (IACUC)
- Consent to participate: Not applicable
- Consent for publication: Not applicable
- Availability of data and materials: All data and materials are available upon request.
- Competing interests: The authors declare no competing interests

## AUTHOR CONTRIBUTIONS

MMA and SO conceptualized, designed, and supervised this study. DS developed methodologies and conducted data acquisition. DS, SS, SO, & MMA carried out the analysis and interpretation of data. JMV, RZ, SO & MMA provided administrative, technical, or material support. DS, JMV, RZ, SO, & MMA conducted writing, reviewing, and/or revising the manuscript.

## Data Availability

The MERFISH data in this study were deposited in the Gene Expression Omnibus (GEO) repository under accession code GSE322277, https://www.ncbi.nlm.nih.gov/geo/query/acc.cgi?acc=GSE%20322277.

The bulk RNA seq data were deposited in the GEO repository under accession number GSE322681, https://www.ncbi.nlm.nih.gov/geo/query/acc.cgi?acc=GSE322681

